# Protein domain characterization reveals human MIC60 tolerates loss of helical bundle domain

**DOI:** 10.64898/2026.03.19.713006

**Authors:** Stephanie Rockfield, Krishnan Venkataraman, Chih-Hsuan Wu, Randall Wakefield, Alan Wu, Amit Budhraja, Ricardo Rodriguez-Enriquez, Ali Khalighifar, Camenzind G. Robinson, Cai Li, Alexandre F. Carisey, Joseph T. Opferman

## Abstract

The mitochondrial contact site and cristae organizing system (MICOS) is essential for cristae junction formation and inner mitochondrial membrane architecture. To define how MICOS integrity is established and maintained, we generated conditional deletion models of *Immt* (encoding MIC60), a core MICOS subunit, in tissue-specific settings and in cultured cells. Liver-specific deletion of *Immt* in mice induced profound defects in mitochondrial ultrastructure and function, establishing MIC60 as essential for mitochondrial integrity. Notably, despite the severity of the defects, we did not detect increased apoptosis in liver tissue or in cells. To directly link MIC60 structure to its function, we performed a systemic structure-function analysis of human MIC60 using domain-specific deletion mutants expressed in *Immt*-deleted cells. We identified that the transmembrane, coiled-coil, and mitofilin domains are required for MICOS assembly, mitochondrial morphology, and respiratory function. Unexpectedly, deletion of the predicted helical bundle (a region spanning 229 amino acids) substantially restored mitochondrial structure and function, nearly matching full-length MIC60. A mutation (K299E) associated with human disease within this domain similarly preserved most MIC60-dependent functions. Together, these results establish MIC60 as a non-redundant regulator of mitochondrial architecture while revealing that a large predicted structural domain is largely dispensable for MIC60’s core functions, refining current models of MICOS organization and uncovering unexpected modularity within MIC60.

## INTRODUCTION

Mitochondria are responsible for many aspects of cellular health, including ATP synthesis, lipid oxidation, and apoptotic regulation^1^. Mitochondrial structures include the outer mitochondrial membrane (OMM) and the inner mitochondrial membrane (IMM), which folds into crista^2–4^ and surrounds the matrix, where mitochondrial DNA (mtDNA) is stored^3^. Crista junctions house the large, multi-protein complex known as the mitochondrial contact site and cristae organizing system (MICOS), which is important for the formation and morphology of the mitochondrial cristae on the IMM^5^. The sorting and assembly machinery (SAM) complex rests within the OMM and coordinates with the transfer of outer membrane (TOM) complex to translocate proteins from the cytoplasm into the mitochondria as well as integrate large β-barrel proteins within the OMM^6^. MICOS proteins MIC60 (gene name *IMMT*) and MIC19 (gene name *CHCHD3*) span across the intermembrane space (IMS) to form the mitochondrial intermembrane bridge (MIB) with the SAM complex^5, 7^.

Prior work in cultured cells has shown that reduced MIC60 expression leads to disrupted cristae morphology and impaired mitochondrial respiration, establishing MIC60 as a core protein within the MICOS complex^8–13^. Multiple studies have found that germline *Immt* deletion results in embryonic lethality in mice^14, 15^. Furthermore, inducible, whole-body deletion of *Immt* in adult mice caused rapid mortality, corresponding with reduced bone marrow cellularity and impaired intestinal function^16^. However, these prior studies have failed to comprehensively assess the impact of MIC60 ablation on MICOS function *in vivo.* Furthermore, while recent work in yeast has begun to elucidate the structure of MIC60^17^, the full structure has yet to be solved. Thus, our understanding of the MIC60 protein domains stems primarily from truncation mutations expressed in yeast or mammalian cells. The N-terminal mitochondrial targeting sequence is cleaved when MIC60 is imported into the mitochondria^18, 19^. The next predicted structured region is the transmembrane domain (TM)^9^, which is predicted to anchor MIC60 to the IMM^19^ and has been reported to be essential for MIC60 function in yeast^20^. The TM domain is followed by a predicted intrinsically disordered region^21^ which has been hypothesized to act as a diffusion barrier between the IMS and the crista lumen^17^. A helical bundle (HB) comprised of multiple alpha-helices is conserved in MIC60 from animals but is absent in yeast^7^. Although the function of this domain has not yet been determined^22^, there are reports of a pathological 895A>G point mutation (Lys299 mutating to Glu, hereafter referred to as K299E) within this domain associated with developmental encephalopathy and optic neuropathy in human patients^23^. The crystal structure for the yeast coiled-coil (CC) domain^18^ has been solved and suggested that MIC60 may organize into tetramers^17^. While some studies have shown that removing the CC domain impaired MICOS organization^24^ and cristae junction formation^20^, others have shown that deleting the CC domain only moderately impacted growth and did not alter MICOS expression in yeast^10^. The CC domain was required for *in vitro* binding of human MIC60 (hMIC60) to MIC19^12^, but its functional importance in mammalian cells remains unexplored. Lipid binding sites and a highly conserved mitofilin domain (MD) are located at the C-terminus of MIC60. Research in yeast has shown that the MD is essential for MIC60 function^10, 20^, as the MICOS protein MIC19 interacts with MIC60 at the MD, exposing the lipid binding sites so they can interact with and bend the mitochondrial membrane^17, 25^. *In vitro* research with recombinant *Drosophila melanogaster* domain mutants revealed that MIC60 interacts directly with the mitophagy regulator PINK1 through the MD^26^. In another study, researchers expressed large C-terminal truncation mutants of hMIC60 in mouse embryonic fibroblasts (MEFs) with shRNA targeting *Immt*^12^ and found that regions spanning both the CC and MD were important for MIC60 function. Despite this extensive investigation of the function of MIC60 protein domains in yeast or mammalian cells, a comprehensive analysis of hMIC60 protein domains and their roles in MICOS function remain under-explored.

Herein, we utilized our conditional *Immt* mouse model^16^ to genetically ablate *Immt* in a tissue-specific manner. We report that loss of MIC60 protein expression in the liver is lethal, correlating with reduced MICOS/MIB protein levels and enlarged mitochondria with aberrant cristae morphology. Additionally, we generated MEFs from our conditional mouse model to facilitate tamoxifen-inducible *Immt* deletion and found similar defects in MICOS protein expression and mitochondrial morphology. Re-expressing different human *IMMT* (h*IMMT*) mutants in these conditional MEFs showed that losing the TM, CC, or MD domains was detrimental to MIC60 interactions with MICOS family members, resulting in aberrant mitochondrial morphology and reduced function. Despite spanning 229 amino acids, removing the HB domain preserved MICOS interactions while maintaining normal mitochondrial function similarly to full-length hMIC60. We also generated the K299E pathogenic mutation and found that expressing this mutant form of hMIC60 in MEFs rescued the effects of *Immt* deletion, although mtDNA organization was impacted. Intriguingly, both the ΔHB and K299E mutants appeared to have a more fragmented mitochondrial network relative to the full-length hMIC60 rescue, suggesting a minor impairment in morphology that overall did not impair mitochondrial function. Together, these results demonstrate the essential function that the majority of the MIC60 protein domains serve towards its function in mammals while identifying a surprisingly dispensable alpha-helical region that expands upon our understanding of MICOS and mitochondrial health.

## RESULTS

### *Immt* deletion in liver is lethal, causing liver damage and aberrant mitochondrial morphology

To determine the impact of tissue-specific ablation of MIC60 in adult mouse livers, control (wild-type, WT) or *Immt*^f/f^ mice were injected with adeno-associated virus (AAV) via tail-vein injections to express Cre recombinase driven by a liver specific promoter (LP1-Cre AAV)^27^. Western blot analysis from liver lysates collected between two- and five-weeks post-injection (p.i.) demonstrated efficient loss of MIC60 within 3 weeks (**Supplemental Figure 1a**). This loss was associated with reduced expression of other MICOS/MIB family members (e.g., MIC19, MIC10, and SAM50) without affecting expression of other mitochondrial proteins such as TOM20 (**Figure 1a**). Importantly, MIC60 expression was unaffected in the spleen, heart, or skeletal muscle following LP1-Cre AAV (**Supplemental Figure 1b**), demonstrating the tissue specificity of this approach.

**Figure 1.**
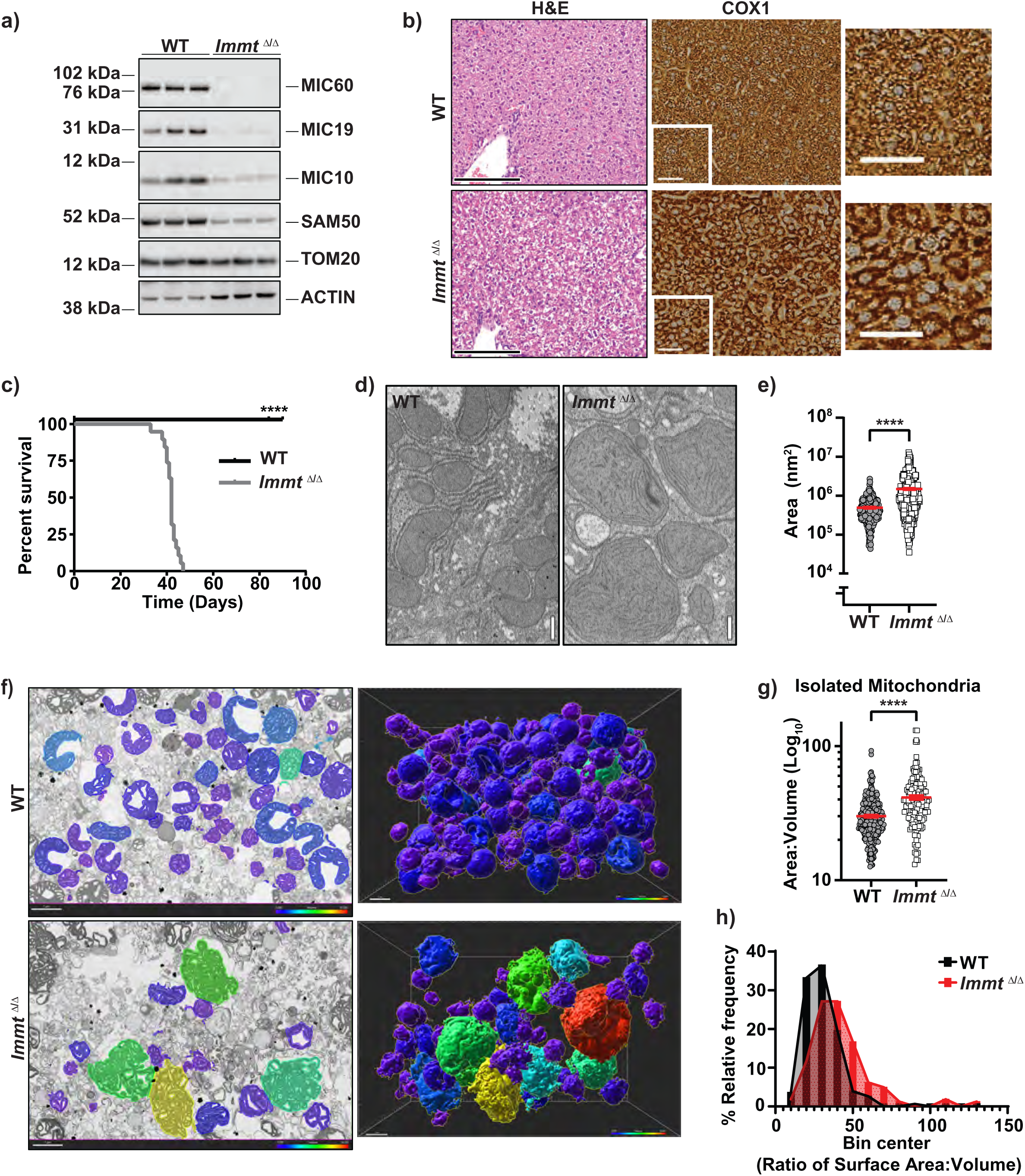
*Immt* deletion in liver is lethal, causing liver damage and aberrant mitochondrial morphology. **a)** Western blot analysis of whole liver lysates 3 weeks after LP1-Cre injection (n = 3 mice/group, representative of 3 independent experiments). **b)** Representative histology images of whole liver lysates 3 weeks after LP1-Cre injection from wild-type (WT; n = 4 M, 3 F) and *Immt*^Δ/Δ^ (n = 4 M, 5 F) mice. For hematoxylin & eosin (H&E) images, scale bar = 100 μm. For COX1 images, scale bar = 50 μm. **c)** Mantel-Cox survival analysis of WT (n = 6 M, 5 F) and *Immt*^Δ/Δ^ (n=6 M, 13 F) mice after LP1-Cre injection. **d)** Representative electron micrographs of whole liver tissue 3 weeks after LP1-Cre injection from WT (n=4 M, 1 F) and *Immt*^Δ/Δ^ (Δ/Δ; n=4 M, 3 F), scale bar = 0.5 μm. **e)** Mitochondrial area of the liver mitochondria shown in (d). Statistical significance determined by an unpaired Welch’s *t*-test. **f)** Electron micrograph (left) and 3D volume (from focused ion beam electron microscopy, right) of isolated liver mitochondria 3 weeks after LP1-Cre injection. Scale bar = 1 μm. Mitochondria are colored according to their volume within a 3D space, with blue being smallest and red being largest. **g)** Area-to-volume ratio from the isolated mitochondria shown in (f). Two independent stacks for each genotype are included (WT n=307 mitochondria, *Immt*^Δ/Δ^ n=162 mitochondria). Statistical significance was determined by an unpaired Welch’s *t*-test. **h)** Relative frequency of mitochondria from (f) falling within the center of an Area:Volume bin. All error bars represent the standard error of the mean. **** *p* ≤ 0.0001.

To assess the effect of MIC60 ablation in livers following LP1-Cre AAV injection, we used immunohistochemical staining for the mitochondrial protein COX1 on liver tissue. Although no marked changes in the expression levels of electron transport chain markers were observed (**Supplemental Figure 1c**), immunohistochemistry detected mitochondrial aggregation in *Immt*^Δ/Δ^ livers, but not WT livers, 3-weeks p.i. (**Figure 1b**). *Immt*^Δ/Δ^ mice died as early as 33 days (4.7 weeks) p.i., with a median survival of 42 days, while WT mice remained unaffected through the duration of the study (**Figure 1c**). Although there was no evidence of apoptosis in the liver either through the presence of cleaved caspase-3 over time (**Supplemental Figure 1a**) or through tissue histology of cleaved caspase-3 and TUNEL (**Supplemental Figure 1d**), we found significantly increased levels of liver damage markers alanine transaminase (ALT) and aspartate aminotransferase (AST)^28^ in blood sera from *Immt*^Δ/Δ^ mice, but not WT mice (**Supplemental Figure 1e**). This increase coincided with the observed negative impact on survival (**Figure 1c**). Since *IMMT* knockdown promotes cytochrome *c* release in HeLa cells^29^, we isolated mitochondria from WT, *Immt*^Δ/WT^, and *Immt*^Δ/Δ^ livers and exposed them to the BID–stapled alpha-helical peptide (SAHB) to assess mitochondrial outer membrane permeability (MOMP)^30, 31^. We found that *Immt*^Δ/Δ^ mitochondria were more primed for MOMP, releasing cytochrome *c* with exposures as low as 10 nM of BID-SAHB as opposed to the 100 nM concentration required for release in the WT and *Immt*^Δ/WT^ samples (**Supplemental Figure 1f**). Together, these results demonstrate that MIC60 is essential for liver function in adult mice, although its loss *in vivo* was not associated with increased apoptosis.

Previous research has established that losing MIC60 expression results in an aberrant mitochondrial morphology^8–13, 16^. To investigate the mitochondrial ultrastructure following *Immt* deletion, we assessed scanning electron transmission microscopy (STEM) scans of whole liver tissue and found that the mitochondria from *Immt*^Δ/Δ^ livers were significantly larger than WT controls (**Figure 1d, e**). Focused ion beam electron microscopy (FIBSEM) micrographs of isolated liver mitochondria also show that *Immt*^Δ/Δ^ livers had significantly larger mitochondria relative to WT mitochondria (**Figure 1f–h, Supplemental Videos 1** and **2**). When comparing the STEM and FIBSEM scans, we observed a clear difference in the cristae morphology in *Immt*-deleted livers that was not apparent in WT controls. Altogether, these results show that liver-specific *Immt* deletion results in liver damage and reduced survival while also affecting mitochondrial size and cristae morphology, providing additional *in vivo* evidence for MIC60’s role in mitochondrial function and overall organismal health.

### *Immt* deletion *in vitro* recapitulates mitochondrial defects observed *in vivo*

To understand the impact of *Immt* deletion on mitochondrial morphology and function, conditional (tamoxifen-inducible) *Immt*-knockout MEFs from the Rosa-ERT^2^ Cre *Immt*^f/f^ mice were established in which treatment with 4-OH-tamoxifen (TAM) could induce Cre-recombinase activity. Addition of TAM to cultured cells reduced MIC60 expression within 2 days and MIC60 expression was nearly ablated by day 4 of TAM treatment (**Supplemental Figure 2a**). The loss of MIC60 also triggered a reduction in the expression of other MICOS/MIB family members without affecting TOM20 expression (**Figure 2a** and **Supplemental Figure 2a**). Importantly, MICOS protein levels were not impacted by TAM in either *Immt*^Δ/WT^ MEFs (**Supplemental Figure 2a**) or WT MEFs (**Figure 2a**).

**Figure 2.**
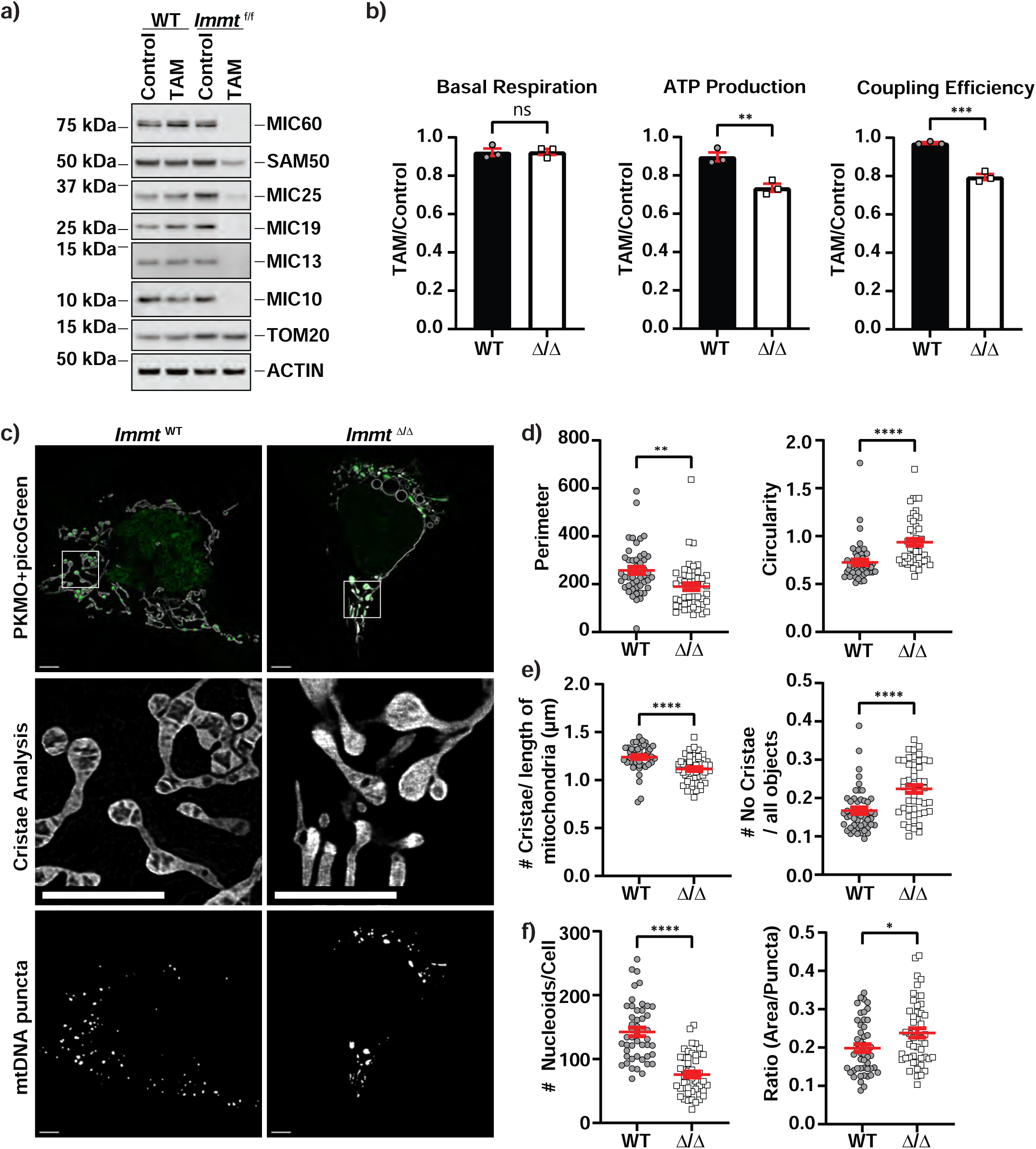
*Immt* deletion *in vitro* recapitulates the mitochondrial defects observed *in vivo.* **a)** Western blot analysis of mouse embryonic fibroblasts (MEFs) generated from wild-type (WT) or *Immt*^f/f^ embryos and treated for 4 days with DMSO (control) or 4-OH-tamoxifen (TAM). Data is representative of 3 independent experiments. **b)** Agilent Seahorse analysis for mitochondrial oxygen consumption in WT (n=3) and *Immt*^Δ/Δ^ (Δ/Δ, n=3) MEFs 4 days post-treatment. Results depict the fold change of TAM to control (DMSO) and represent 3 independent experiments. **c)** Representative live cell tauSTED images of MEFs at 4 days of TAM treatment. Cells were stained with picoGreen (green) and PKMO (gray). Scale bar = 5 μm. n = 48 cells each. Images are representative of 3 independent experiments. **d)** Differences in mitochondrial morphology (perimeter (in pixels), left, and circularity, right) in WT and *Immt*^Δ/Δ^ MEFs shown in (c) at 4 days of TAM treatment. **e)** Differences in density of mitochondrial cristae (left) and number of mitochondrial objects containing no discernable cristae (right) in WT and *Immt*^Δ/Δ^ MEFs shown in (c) at 4 days of TAM treatment. **f)** Differences in quantity (left) and average area (right) of mitochondrial nucleoids (puncta) in WT and *Immt*^Δ/Δ^ MEFs shown in (c) at 4 days of TAM treatment. Statistical significance was determined by unpaired Welch’s *t*-test. Data represent mean ± SEM from 3 independent experiments. * *p* ≤0.05, ** *p* ≤ 0.01, *** *p* ≤0.001, **** *p* ≤ 0.0001, ns = not significant.

To ascertain the specific impact of MIC60 loss on MEF mitochondrial function, we assessed respiration after *Immt* deletion (**Figure 2b** and **Supplemental Figure 2b and c**). Although we did not find changes in basal or maximal respiration, we observed a trend towards reduced spare respiratory capacity and significantly reduced ATP levels, reduced electron coupling efficiency, and elevated proton leak (an indicator of damaged mitochondria^32^) between days 4 and 5 of TAM treatment. While cell proliferation was significantly hindered in *Immt*-deleted MEFs after 3 days of TAM treatment (**Supplemental Figure 2d**), cell viability was not changed even by day 7 (**Supplemental Figure 2e**). Furthermore, we did not see evidence of caspase-3 cleavage (**Supplemental Figure 2f**), indicating apoptosis was not occurring in our cells after *Immt* deletion. To determine the time course for which induced *Immt* deletion affects mitochondrial morphology, we used structured illumination microscopy (SIM) super-resolution microscopy of the inducible knockout MEFs. We found evidence of rounded mitochondria as early as day 2 of TAM treatment and enlarged, rounded mitochondria prevalent by day 4 (**Supplemental Figure 3a**). We also used the STimulated Emission Depletion (STED)-able mitochondrial dye PKmito ORANGE (PKMO), enabling us to visualize mitochondria and cristae morphology in live cells^33^. Representative 2D images from WT and *Immt*^Δ/Δ^ MEFs are shown in **Figure 2c** (top) and revealed a striking difference in mitochondrial morphology. We took an unbiased approach to quantify mitochondrial features including area, axis length, perimeter, circularity, and texture. Statistical analysis from a linear mixed model (LMM) revealed a significant difference (p=0.029) in solidity, circularity, area-to-perimeter ratio, major axis length, and minor axis length (**Supplemental Table 1**); perimeter and circularity are presented in **Figure 2d**. Next, WT and *Immt*^Δ/Δ^ cells were manually assessed as previously described^12^, scoring cells by the presence of elongated and branched, fragmented, or enlarged and rounded mitochondria. We found that only ∼4% of WT MEFs had mitochondria that could be classified as enlarged and rounded whereas ∼92% of *Immt*^Δ/Δ^ MEFs contained enlarged and rounded mitochondria (**Supplemental Table 2**).

We next investigated whether *Immt* deletion affected the cristae network, using WT and *Immt*^Δ/Δ^ MEFs stained with PKMO. We found significantly fewer cristae per length of mitochondria with *Immt* deletion, as well as significantly increased prevalence of mitochondria lacking discernable cristae in *Immt*^Δ/Δ^ MEFs (**Figure 2e**). Cells were co-stained with the double-strand DNA dye picoGreen^34^ so we could characterize mtDNA nucleoid organization. Puncta analysis of mtDNA revealed significantly fewer nucleoids per cell with *Immt* deletion, although the nucleoids were significantly larger compared to the WT MEFs (**Figure 2f**). We next used tetramethylrhodamine (TMRM) dye and confocal microscopy to determine whether *Immt* deletion impacted mitochondrial membrane potential (ψ). As shown in **Supplemental Figure 3b and c**, we found significantly reduced ψ following *Immt* deletion relative to WT. Overall, our inducible cell culture system demonstrated that *Immt* loss coincided with aberrant mitochondrial morphology, disrupted cristae and mtDNA networks, reduced ψ, and impaired mitochondrial function resulting from reduced MIC60 expression.

In 63% (30/48 cells) of *Immt*^Δ/Δ^ MEFs, we observed the presence of large, spherical mitochondria that were devoid of membrane or mtDNA nucleoids (**Figure 2c**); we did not see this phenomenon in our fixed-cell SIM images, although we saw rounded mitochondria with a ring of TOM20-stained OMM filled with cytochrome *c* (**Supplemental Figure 3a**, day 5). Since mtDNA is localized to the mitochondrial matrix while cytochrome *c* is within the IMS^4, 35^, we hypothesized that these mitochondria were comprised primarily of IMS that was not being detected with our PKMO and picoGreen dyes. To test this, we stably expressed IMS-RP^36^ in *Immt*^f/f^ MEFs and completed live-cell confocal imaging using PKmito Deep Red (PKMDR^37^) and picoGreen (**Supplemental Figure d** and **e**). Control WT (DMSO treated, top images) showed an overall elongated morphology with even distribution of IMS-RP and interchanging intensities between the PKMDR and picoGreen. The spherical mitochondria seen after *Immt* deletion (bottom images) were comprised of either IMS-RP or mtDNA, a marked shift from the morphology of WT MEFs.

### hMIC60 protein domain mutants reveal helical bundle is dispensable for MICOS

We next set out to complete a structure-function analysis of hMIC60 using our inducible *Immt* knockout system; the AlphaFold2 predicted structure is presented in **Figure 3a**. We generated deletion mutants for the TM domain (ΔTM, amino acids [aa] 41–62), the HB (ΔHB, aa 203–431), the CC helix (ΔCC, aa 432–697), and the evolutionarily conserved MD (ΔMD, aa 698–758) all with a C-terminal V5 epitope tag. These V5-tagged constructs, as well as the empty vector (EV) and full-length h*IMMT* controls, were stably expressed in *Immt*^f/f^ MEFs. We demonstrated that h*IMMT* rescued the expression levels of MICOS and MIB proteins although the faster migrating endogenous MIC60 was absent (**Figure 3b**). Neither ΔTM, ΔCC, nor ΔMD constructs were able to rescue the loss of MICOS protein levels in TAM-treated cells (we note here that the anti-MIC60 antibody epitope is at the C-terminus and thus does not react to the ΔMD mutant). In contrast, the ΔHB mutant fully rescued MICOS/MIB protein levels as well as full-length hMIC60, an unexpected result given its size and specific presence in animals^7, 22^.

**Figure 3.**
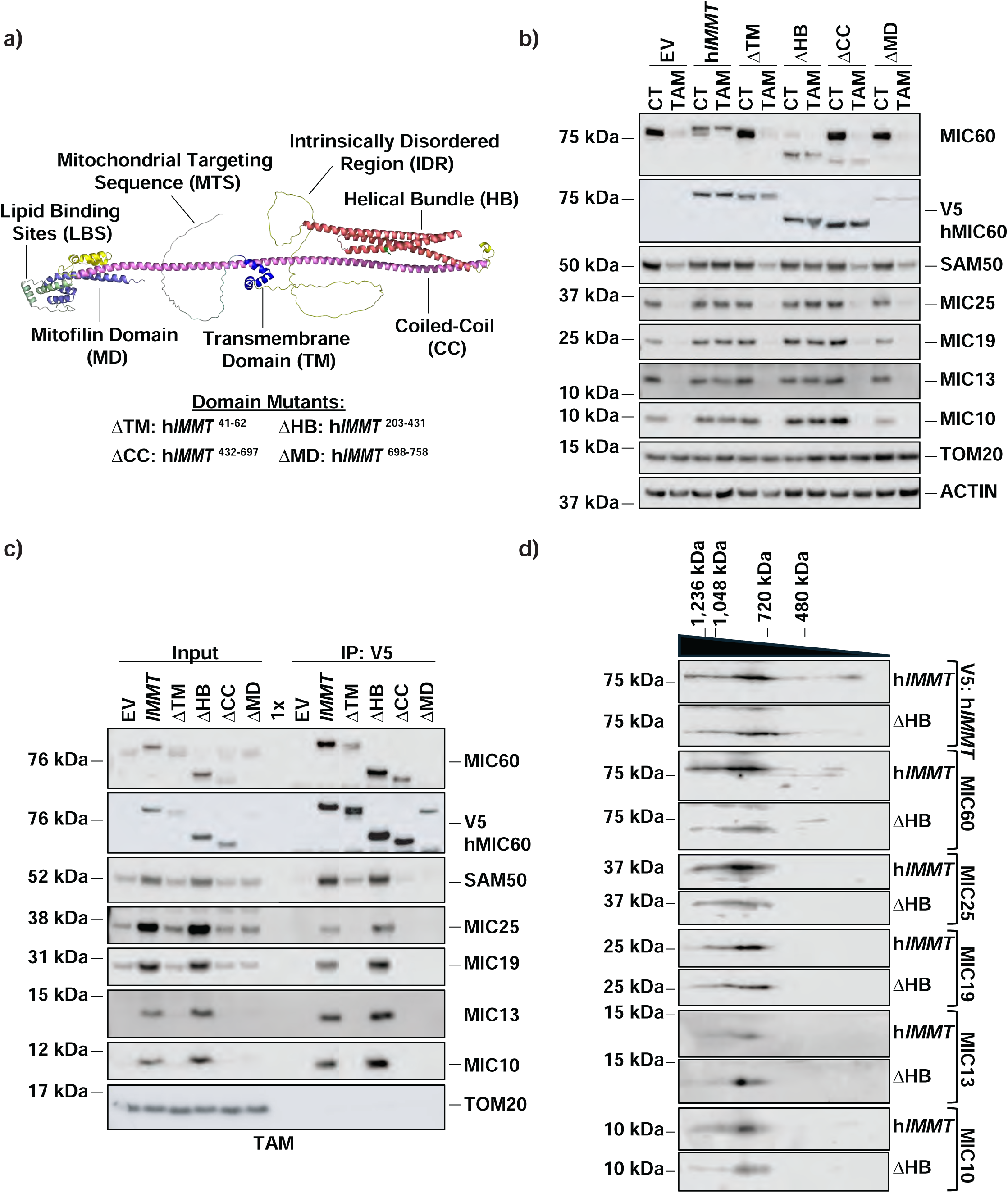
hMIC60 protein domain mutants reveal helical bundle is dispensable for MICOS. **a)** AlphaFold2 predicted structure of hMIC60 (gene name: *IMMT*). Predicted protein domains are colored as follows: mitochondrial targeting sequence (gray), transmembrane domain (TM, dark blue), intrinsically disordered region (yellow), helical bundle (HB, coral), coiled-coil (CC, pink), lipid binding sites (light green), and mitofilin domain (MD, violet). The K299 residue is indicated within the HB domain in green. The protein domain mutants used in this study are summarized at the bottom of this panel. **b)** Western blot analysis of mouse embryonic fibroblasts (MEFs) stably expressing V5-tagged h*IMMT* and protein domain deletion mutants treated for 4 days with DMSO (control, CT) or 4-OH-tamoxifen (TAM; n = 3 independent experiments). **c)** Western blot analysis of immunoprecipitated proteins pulled down with V5 after 4 days TAM treatment (n = 3 independent experiments). **d)** Two-dimensional blue-native PAGE of isolated mitochondria from h*IMMT* or ΔHB mutant after 4 days TAM treatment (n = 3 independent experiments). Abbreviation: EV, empty vector.

We tested whether these hMIC60 mutants could interact with the endogenous MICOS and MIB family members. Both exogenous hMIC60 and ΔHB fully interacted with all assessed MICOS and MIB proteins (**Figure 3c** and **Supplemental Figure 4a**). Even with the presence of endogenous MIC60 (no TAM), the ΔTM, ΔCC, and ΔMD mutants did not immunoprecipitate with the MICOS proteins (despite both ΔCC and ΔMD mutants interacting with murine MIC60; **Supplemental Figure 4a**). When treated with TAM, the ΔHB mutant immunoprecipitated the MICOS family proteins as efficiently as the full-length hMIC60 (**Figure 3c**). We also completed the MICOS assembly assay^38^, isolating mitochondria from TAM-treated h*IMMT* and ΔHB-expressing MEFs followed by 2D-blue native PAGE. As shown in **Figure 3d**, MICOS assembly was equivalent in both cell lines, further demonstrating that deletion of the large HB domain did not impair MICOS complex assembly or interactions.

We next investigated the impact of these mutants on mitochondrial function and morphology. Exogenous expression of full-length hMIC60 rescued basal respiration, maximal respiration, and ATP production (**Figure 4a**). Similarly, the ΔHB mutant significantly rescued basal respiration, maximal respiration, and ATP production. Expression of either ΔTM, ΔCC, or ΔMD failed to rescue any of the assessed respiratory functions (**Figure 4a**).

**Figure 4.**
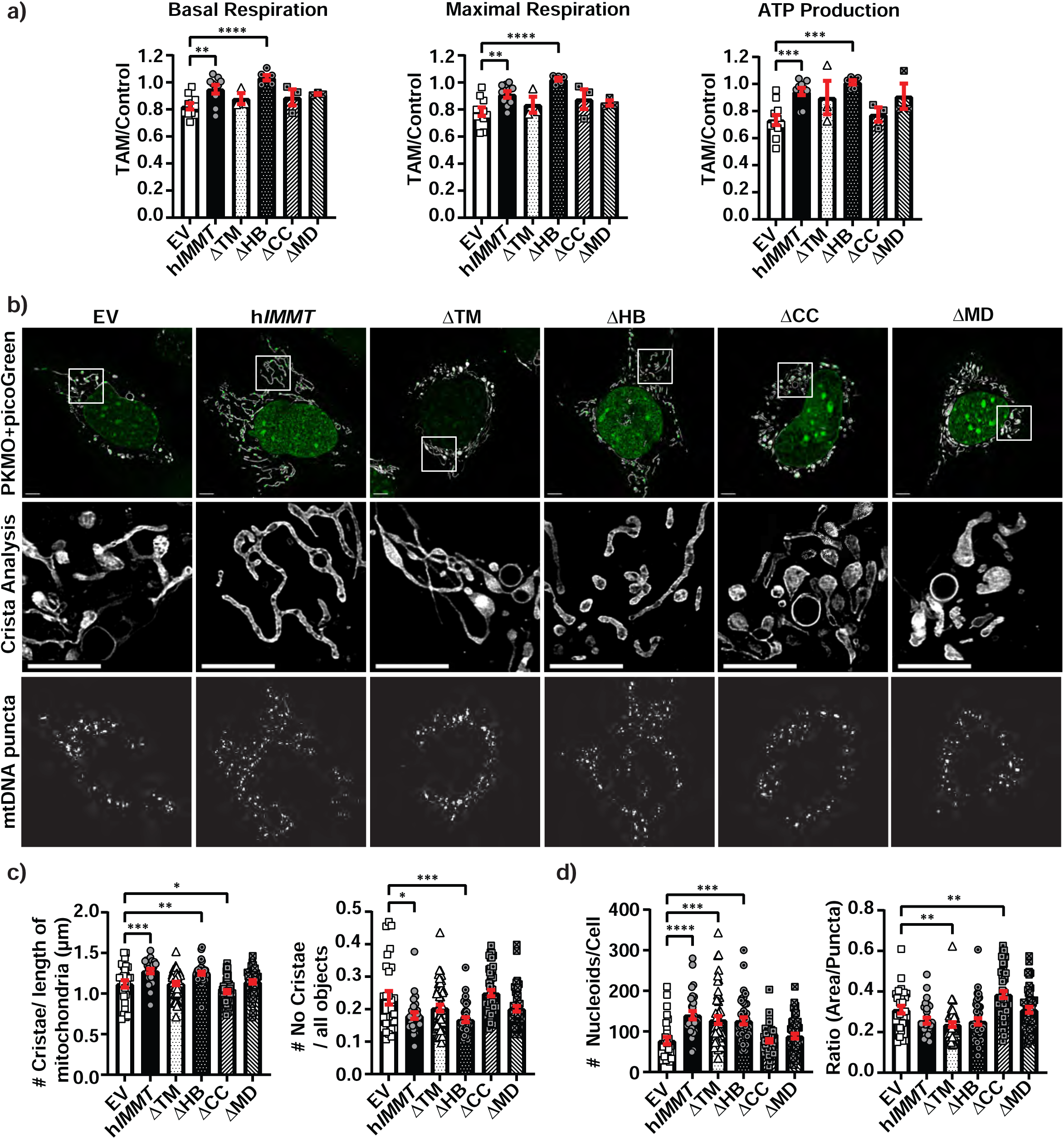
hMIC60 protein domain mutants reveal helical bundle is dispensable for mitochondrial function but only partially rescues morphology. **a)** Agilent Seahorse analysis for mitochondrial oxygen consumption in mouse embryonic fibroblasts (MEFs) expressing V5-tagged h*IMMT* and protein domain deletion mutants (transmembrane domain [ΔTM], helical bundle [ΔHB], the coiled-coil helix [ΔCC], and mitofilin domain [ΔMD]) 4 days 4-OH-tamoxifen (TAM) post-treatment. Results depict the fold change of TAM to control (DMSO) and represent at least 3 independent experiments. **b)** Representative live cell tauSTED images of h*IMMT* mutant-expressing MEFs at 4 days of TAM treatment. Cells were stained with picoGreen (green) and PKMO (gray). Scale bar = 5 μm. Cells per group are as follows: empty vector (EV) = 29, h*IMMT* = 29, ΔTM = 50, ΔHB = 39, ΔCC = 50, ΔMD = 50. **c)** Differences in density of mitochondrial cristae (left) and number of mitochondrial objects containing no discernable cristae (right) in MEFs expressing h*IMMT* and its mutants shown in (b) at 4 days of TAM treatment. **d)** Differences in quantity (left) and average area (right) of mitochondrial nucleoids (puncta) in MEFs expressing h*IMMT* and its mutants shown in (b) at 4 days of TAM treatment. Data in (a, c, d) are presented as mean ± SEM from 3 independent experiments. Statistical significance was determined using one-way ANOVAs; * *p* ≤0.05, ** *p* ≤ 0.01, *** *p* ≤ 0.001, and **** *p* ≤ 0.0001.

To assess whether exogenous expression of MIC60 mutants was able to restore normal mitochondrial morphology, we visualized reconstituted MEFs using live-cell STED. As shown in **Figure 4b**, the EV as well as the ΔTM, ΔCC, and ΔMD mutants all displayed abnormal, enlarged, rounded mitochondria characteristic of *Immt* deletion while the expression of h*IMMT* resulted in a normal mitochondria network featuring elongated and branched mitochondria with clearly discernable cristae. Although our LMM did not identify any significant differences between the mutants (**Supplemental Table 3**), we found that between 80%–90% of EV, ΔTM, ΔCC, and ΔMD mutant MEFs contained enlarged and rounded mitochondria compared to ∼3% of h*IMMT* MEFs (**Supplemental Table 4**). Intriguingly, we found that the ΔHB mitochondria were not as elongated as the h*IMMT-*expressing mitochondria even though this mutant prevented the generation of enlarged, rounded mitochondria. Our manual scoring of the images found that ∼13% of the ΔHB MEFs contained enlarged and rounded mitochondria, with ∼67% of the MEFs containing fragmented mitochondrial morphologies (compared to ∼55% of h*IMMT* cells, **Supplemental Table 4**). Despite this change in mitochondrial morphology, we found that both cristae density and the number of mitochondria containing no discernable cristae were equally rescued with h*IMMT* or ΔHB (**Figure 4c**). In addition, ΔCC-expressing MEFs showed significantly reduced cristae density (**Figure 4c**), suggesting a more drastic phenotype compared to the EV (*Immt*-deleted) MEFs. When assessing mtDNA nucleoids, we found that h*IMMT* had significantly increased the number of mtDNA nucleoids that trended towards being smaller relative to the EV controls, and the ΔHB-expressing MEFs were comprised of mitochondrial nucleoids similar in size and quantity to the h*IMMT* MEFs (**Figure 4d**). While ΔTM-expressing MEFs contained large nucleoids morphologically similar to MEFs with *Immt* deletion, they, overall, showed significantly increased nucleoid numbers that were smaller in size. Furthermore, the average nucleoid size for ΔCC-expressing MEFs was significantly increased relative to EV (**Figure 4d**). Altogether, these results indicate that the TM, CC, and MD domains are all essential for promoting MIC60 function and preserving mitochondrial morphology, while the HB domain appears dispensable. The ΔHB mutant rescued MICOS expression and interactions while restoring cristae morphology and mitochondrial function, although the ΔHB-expressing MEFs were comprised of a more fragmented mitochondrial network compared to the MEFs expressing full-length hMIC60.

### Characterization of the pathogenic K299E mutant

We next investigated the known pathogenic mutation, K299E^23^, in h*IMMT*. Expression of the K299E mutant rescued MICOS/MIB protein levels similarly to the full-length hMIC60 rescue previously observed (**Figure 5a**). Using immunoprecipitation, we found that the K299E mutant retained interactions with the murine MICOS proteins in both DMSO control (**Supplemental Figure 4b**) and TAM-treated conditions (**Figure 5b**). We also found that electron coupling efficiency was significantly rescued relative to the EV *Immt-*deleted MEFs while there was a trend towards improved ATP production (**Figure 5c**). These results suggest that the K299E mutation did not negatively affect MICOS complex assembly or mitochondrial function.

**Figure 5.**
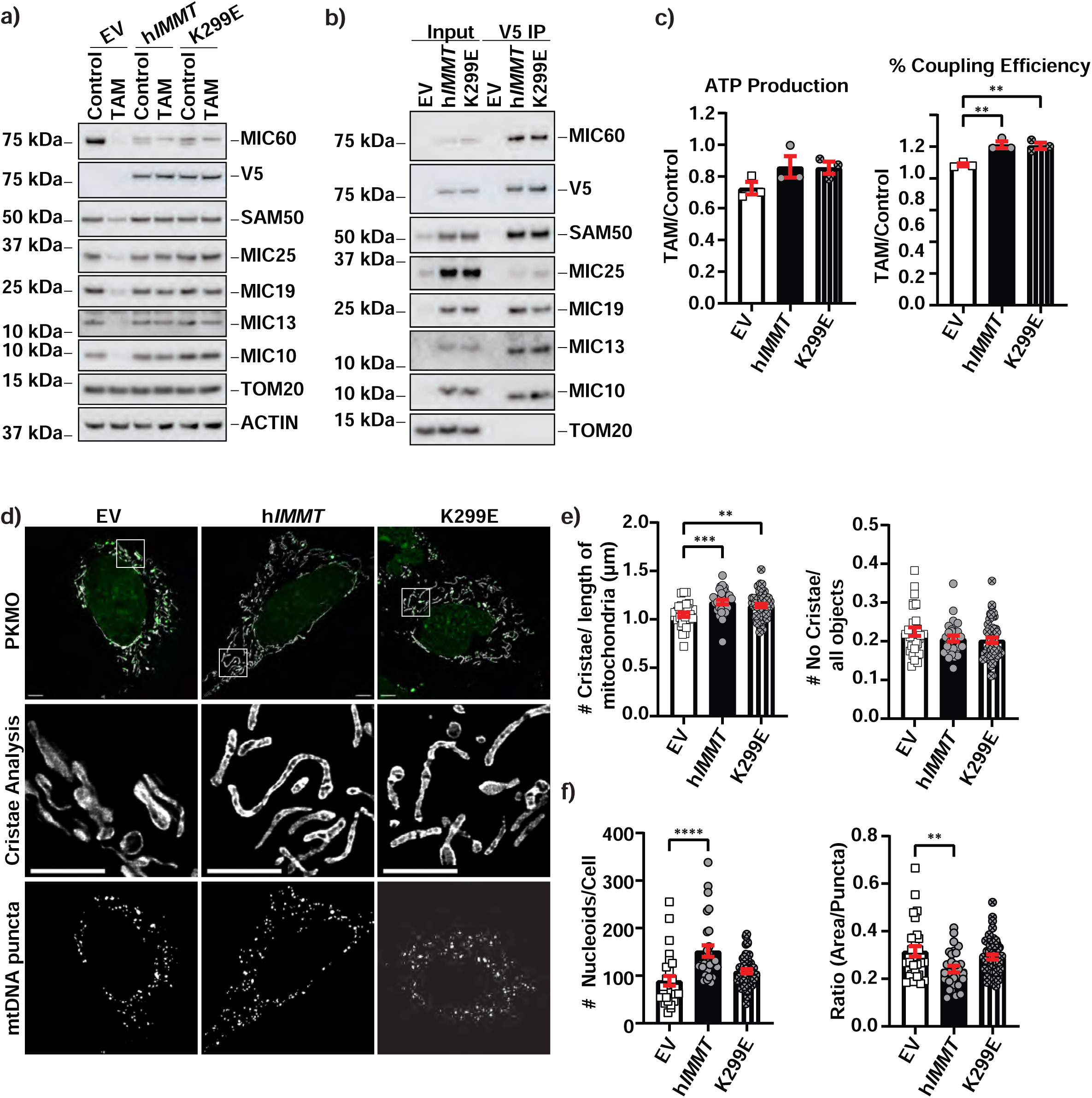
Characterization of pathogenic K299E mutant. **a)** Western blot analysis of mouse embryonic fibroblasts (MEFs) stably expressing an empty vector (EV), h*IMMT*, or the K299E mutant after being treated with DMSO (control) or 4-OH-tamoxifen (TAM) for 4 days (n = 3 independent experiments). **b)** Western blot analysis of immunoprecipitated proteins pulled down with V5 after TAM treatment for 4 days (n = 3 independent experiments). **c)** Agilent Seahorse analysis for mitochondrial oxygen consumption in MEFs expressing EV, h*IMMT*, or the K299E mutant 4 days post TAM treatment. Results depict the fold change of TAM to control (DMSO) and represent 3 independent experiments. **d)** Representative live-cell tauSTED images of MEFs expressing EV (n = 29), h*IMMT* (n = 29), or K299E mutant (n = 50) at 4 days of TAM treatment. Cells were stained with picoGreen (green) and PKMO (gray). Scale bar = 5 μm. **e)** Differences in density of mitochondrial cristae (left) and number of mitochondrial objects containing no discernable cristae (right) in MEFs expressing EV, h*IMMT*, or K299E mutant shown in (d) at 4 days of TAM treatment. **f)** Differences in quantity (left) and average area (right) of mitochondrial nucleoids (puncta) in MEFs expressing EV, h*IMMT*, or K299E mutant shown in (d) at 4 days of TAM treatment. Data in (c, e, f) presented as mean ± SEM from 3 independent experiments. Statistical significance was determined using one-way ANOVAs; * *p* ≤ 0.05, ** *p* ≤ 0.01, *** *p*≤0.001, and **** *p* ≤ 0.0001.

We investigated whether mitochondrial morphology was impacted by this pathogenic mutation. As shown in **Figure 5d**, K299E MEFs failed to display an enlarged, rounded mitochondrial phenotype, supporting its ability to rescue *Immt* deletion. In fact, our manual scoring method found that only ∼20% of the K299E mutant MEFs contained enlarged and rounded mitochondria, while ∼76% of these MEFs contained a fragmented morphology reminiscent of the ΔHB mutant (**Supplemental Table 4**). We found that mitochondrial cristae density was also rescued with the K299E mutant (**Figure 5e**). Intriguingly, we found that mtDNA nucleoids were more similar to the EV *Immt-*deleted MEFs in both number and size (**Figure 5f**), suggesting this K299E mutation is partially functional yet deficient in some manner that affects mitochondrial morphology and mtDNA organization.

MIC60 is known to interact with the IMS protein OPA1, a mediator of mitochondrial fusion^39–41^ and mtDNA organization^42, 43^. Given that we observed fewer elongated mitochondria in both the ΔHB and K299E mutants (as well as aberrant mtDNA puncta in the K299E mutant), we hypothesized that this protein domain may be important for MIC60’s interaction with OPA1. To map which domains are important for MIC60–OPA1 interactions, we used an immunoprecipitation approach. As shown in **Supplemental Figure 4a**, the TM, CC, and MD domains were all essential for OPA1 pulldown, whereas OPA1 still immunoprecipitated with the ΔHB mutant. EV *Immt*-deleted MEFs had altered OPA1 processing, with fewer long-OPA1 and increased short-OPA1; this effect was completely rescued in h*IMMT*, ΔHB, and K299E MEFs (**Supplemental Figure 4c**). Additionally, upon treatment with TAM, we found that both the ΔHB and K299E mutants retained their interactions with OPA1 (**Supplemental Figure 4d**). We also captured live-cell videos using the PKMO dye and quantified fission and fusion rates using MitoMeter^44^. As shown in **Supplemental Figure 4e**, we did not see a significant change in either fission or fusion in either mutant or even between the EV *Immt-*deleted and h*IMMT* MEFs. These results ultimately suggest OPA1-mediated fusion was not impacted by either the ΔHB or K299E mutant 4 days after *Immt* deletion.

## DISCUSSION

Herein, we comprehensively assessed the contribution of specific structured domains to the functionality of MIC60. By excising these specific regions and expressing in our inducible *Immt* knockout mammalian cells, we definitively demonstrated that the TM, CC, and MD domains are essential for MICOS complex expression (**Figure 3**), as well as mitochondrial morphology and function (**Figure 4**). We found that the HB domain was largely dispensable, as MEFs reconstituted with this mutant retained interactions with other MICOS family members (**Figure 3**), while restoring mitochondrial respiration and preventing the formation of enlarged, rounded mitochondria (**Figure 4**). Given that this is a large domain spanning 229 amino acids is conserved across animal species^7, 22^, the fact that it nearly completely rescues MEFs from the effects of induced *Immt* deletion was unexpected. A recently submitted preprint has provided structural insights into the HB domain from humans, suggesting that MIC60 dimers may interact through these residues^45^. Similarly, a pathogenic variant of *IMMT*^23^, which resides within the HB domain, equally rescued reconstituted MEFs from *Immt* deletion (**Figure 5**). Despite their ability to rescue the observed mitochondrial dysfunction, expression of both the ΔHB and K299E mutants resulted in a more fragmented mitochondrial network, suggesting this domain may have functional roles related to maintaining mitochondria morphology outside of MIC60’s canonical role within MICOS.

Since prior *in vivo* research in rats as well as mammalian cell culture has linked MIC60 expression and apoptosis ^8, 31, 41, 46, 47^, we investigated whether *Immt* deletion was associated with increased cell death in our inducible system. In both adult mouse liver tissue (**Supplemental Figure 1**) and in MEFs (**Supplemental Figure 2**), we did not find evidence of apoptosis induction following *Immt* deletion. Instead, we observed a deficiency in cell growth over time corresponding with the loss of MICOS expression and reduced mitochondrial respiration (**Supplemental Figure 2**). We speculate that unmet energy demands following *Immt* deletion may impact cellular proliferation. In accordance with prior studies in rat myoblast cells^31^, isolated liver mitochondria exhibited enhanced priming for cytochrome *c* release. Our results also show an altered ratio of long- and short-OPA1 isoforms as a result of *Immt* deletion in MEFs (**Supplemental Figure 4**); prior research demonstrates that altered OPA1 processing promoted cytochrome *c* priming^48, 49^, perhaps explaining why MOMP priming was enhanced following the loss of MIC60 expression.

Our assessment of the pathogenic K299E mutation did not identify any overt functional defects in MIC60 function in relation to maintaining MICOS expression or rescuing the enlarged and rounded mitochondrial morphology (**Figure 5**). Unlike MIC13 mutations in young patients presenting with liver dysfunction and developmental delays^50–52^, the *IMMT* K299E mutation was not lethal^23^. This suggests that the mutation may simply impede MIC60 function. Consistent with this impact, we observed a more fragmented mitochondrial network in K299E-expressing MEFs compared to the *Immt*-deleted MEFs rescued with full-length hMIC60 (**Figure 5**). We speculate that assessing this mutation in a more relevant cell type, such as neurons, may reveal more critical deficiencies in MIC60 function caused by the K299E variant.

Our assessment of MIC60’s structured protein domains highlights their functional importance in mitochondrial biology, especially within the regions highly conserved between animals and yeast. Despite being dispensable for mitochondrial function, the HB appears to have functions outside the MICOS complex that remain unclear. In the future, our inducible *Immt*-knockout mouse model (**Figure 1**) could provide key insight in a tissue-specific or even an organismal manner, especially for the pathogenic K299E mutation. Lysine residues are a common target for post-translational modifications^53^. Future research could interrogate whether the K299E impairs post-translational processing at this residue, disrupting a novel MIC60 regulatory mechanism. Indeed, only a few reports have assessed MIC60 signaling regulation to date^26, 54^, representing an underexplored area in the MICOS and mitochondrial regulation fields. By clarifying which MIC60 domains are indispensable and uncovering new roles for the predicted HB region, this study equips the field with a mechanistic roadmap that will guide future efforts to dissect MICOS regulation in health and disease.

### Limitations of study

One limitation of our study is the ectopic expression of hMIC60 within our inducible *Immt*-deletion MEFs, whereby both hMIC60 and mouse MIC60 is expressed up until *Immt* deletion is triggered with tamoxifen. While a genetic knock-in approach may be favored to achieve endogenous expression levels, this would mean that any mutants that failed to rescue MIC60 function would also fail to yield cell lines (since complete loss of *Immt* is lethal^14–16^). Given that MICOS family members are long-lived proteins with slow turnover rates^55, 56^, our system enabled us to genetically ablate endogenous MIC60 expression levels with high consistency relative to transient knockdown techniques. Indeed, it took at least 3 days to see reduced MIC60 protein levels in cultured MEFs (**Supplemental Figure 2**), but this system facilitated our structure/function analysis of these structured protein domains.

Another limitation of our study is that we focused our attention on the liver when targeting *Immt* in a tissue-specific manner (**Figure 1**). As a mitochondria-dense tissue^57^, this was a logical approach to take, especially given the demonstrated ease of using the LP1-Cre AAV system for liver-specific genetic ablation of *Mcl1*^27^. However, this approach prevents us from assessing potential differences in MIC60 function within other tissues. Future work could address this by targeting *Immt* for deletion in other tissues, such as the brain, or even in a cell-type specific manner.

Finally, our analysis of MIC60’s structured protein domains meant that we removed vast regions of the protein spanning over 200 amino acids (**Figure 3**). It is possible that such large truncations may have caused conformational changes that ultimately affected protein function, resulting in an inability to rescue MEFs from *Immt* deletion. However, our inducible system is well-situated for a more precise analysis within these predicted structured domains. Future research could focus on key residues, such as those receiving post-translational modifications^26, 54^, to provide deeper insight into MIC60 signaling regulation.

## Acknowledgements

This work was supported by the American Lebanese Syrian Associated Charities (ALSAC) of St. Jude Children’s Research Hospital (SJCRH). Microscopy work was performed at the CMB/CPNDR Microscopy Core Facility which is supported by SJCRH. STEM and FIBSEM images were acquired at the Cell & Tissue Imaging Center, which is supported by SJCRH and NCI P30 CA021765. The authors are also grateful to the members of the Opferman Laboratory (SJCRH) for helpful discussions. Finally, the authors thank the SJCRH Animal Resource Center, Comparative Pathology Core, Flow Cytometry and Cell Sorting Core, Center for Bioimage Informatics, and Department of Biostatistics for their support of this project.

## STAR★Methods

### Key resources table

| REAGENT or RESOURCE | SOURCE | IDENTIFIER |
| --- | --- | --- |
| Antibodies |  |  |
| MIC60 | Bethyl | Cat#A305-025; RRID: AB_2621219 |
| V5 | Thermo Fisher Scientific | Cat#R960-25, RRID:AB_2556564 |
| SAM50 | Abcam | Cat#ab167430; RRID: AB_3738359 |
| APOOL/MIC27 | Novus | Cat#NBP1-89325, RRID:AB_11015125 |
| CHCHD6/MIC25 | ProteinTech | Cat#20639-1-AP, RRID:AB_10697667 |
| CHCHD3/MIC19 | Thermo Fisher Scientific | Cat#PA5-31578, RRID:AB_2549051 |
| C19orf70/MIC13 | ProteinTech | Cat#25514-1-AP, RRID:AB_2880112 |
| C1orf151/MIC10 | GeneTex | Cat#GTX46630, RRID:AB_11170127 |
| TOM20 | Santa Cruz | Cat#sc-11415, RRID:AB_2207533 |
| Total OXPHOS Rodent WB Antibody Cocktail | Abcam | Cat#ab110413, RRID:AB_2629281 |
| SDHA (Fp70) | Thermo Fisher Scientific | Cat#459200, RRID:AB_10838019 |
| ATP5B | Santa Cruz | Cat#sc-58618, RRID:AB_671743 |
| UQCRC2 | Thermo Fisher Scientific | Cat#PA5-30204, RRID:AB_2547678 |
| MnSOD | BD Biosciences | Cat#611580, RRID:AB_399026 |
| Cytochrome *c* (WB) | Cell Signaling Technology | Cat#11940, RRID:AB_2637071 |
| Cytochrome *c* (IF) | Cell Signaling Technology | Cat#12963, RRID:AB_2637072 |
| OPA1 | BD Biosciences | Cat#612607, RRID:AB_399889 |
| ACTIN | Millipore | Cat#MAB1501, RRID:AB_2223041 |
| GAPDH | Cell Signaling Technology | Cat#2118, RRID:AB_561053 |
| MTCO1/COX1 | Abcam | Cat#ab14705, RRID:AB_2084810 |
| Caspase-3 (WB) | Cell Signaling Technology | Cat#9662, RRID:AB_331439 |
| Cleaved Caspase-3 (IHC) | Cell Signaling Technology | Cat#9664, RRID:AB_2070042 |
| In Situ Cell Death Detection Kit – POD (TUNEL) | Roche | Cat#11684817910, RRID:AB_2861314 |
| Donkey anti-Mouse IgG-HRP | Thermo Fisher Scientific | Cat#SA1-100, RRID:AB_325993 |
| Goat anti-Rabbit IgG-HRP | Bio-Rad | Cat#172-1019,<br>RRID:AB_11125143 |
| Goat anti-Rabbit IgG Alexa Fluor-568 | Thermo Fisher Scientific | Cat#A-11011,<br>RRID:AB_143157 |
| Donkey anti-Mouse IgG Alexa Fluor-568 | Thermo Fisher Scientific | Cat#A10037,<br>RRID:AB_11180865 |
| Donkey anti-Mouse IgG Alexa Fluor-488 | Thermo Fisher Scientific | Cat#A-21202,<br>RRID:AB_141607 |
| Goat anti-Rabbit IgG Alexa Fluor-647 | Thermo Fisher Scientific | Cat#A-21245,<br>RRID:AB_2535813 |
| Donkey anti-Mouse IgG Alexa Fluor-647 | Thermo Fisher Scientific | Cat#A-31571,<br>RRID:AB_162542 |
| Bacterial and virus strains |  |  |
| LP1-Cre adeno-associated virus | St. Jude Vector Core |  |
| pHAGE-UBC-MCS-IZsGreen-W lentivirus | Harvard University | Cat#EvNO00061637 |
| IMS-RP pBABE-puro retrovirus | <sup>36</sup> | Addgene Plasmid<br>24535 |
| Biological samples |  |  |
| Chemicals, peptides, and recombinant proteins |  |  |
| PKmito ORANGE | Cytoskeleton, Inc | Cat#CY-SC053 |
| PKmito DEEP RED | Cytoskeleton, Inc | Cat#CY-SC055 |
| Quant-iT™ PicoGreen™ dsDNA Reagent | Thermo Fisher Scientific | Cat#P11495 |
| MitoTracker™ Green FM | Thermo Fisher Scientific | Cat#M7514 |
| DAPI | Thermo Fisher Scientific | Cat#D1306 |
| Critical commercial assays |  |  |
| Seahorse MitoStress Test Assay | Agilent | N/A |
| Mitochondria Isolation Kit, mouse tissue | Miltenyi | Cat#130-096-946 |
| BCA Protein Assay Kit | Thermo Fisher Scientific | Cat#23227 |
| Deposited data |  |  |
| nnU-Net Training Dataset:<br>https://zenodo.org/records/19071599 | This paper |  |
| nnU-Net Mitochondria Segmentation Model:<br>https://zenodo.org/records/18715522 | This paper |  |
| Experimental models: Cell lines |  |  |
| *Immt*<sup>fl/fl</sup> Rosa ERT2-Cre MEFs | This paper |  |
| Experimental models: Organisms/strains |  |  |
| Mouse: *Immt*<sup>fl/fl</sup>; B6.129-Gt(*ROSA*)26Sor<sup>tm1(cre/ERT2)Tyj/J</sup> | Rockfield *et al.* <sup>16</sup> | N/A |
| Oligonucleotides |  |  |
| See Supplemental Table 5 for cloning primers | This paper | N/A |
| Recombinant DNA |  |  |
| Human *IMMT* | Origene | RC201854L1V |
| ΔTM (h*IMMT*<sup>41-62</sup>) | This paper | |
| ΔHB (h*IMMT*<sup>203-431</sup>) | This paper | |
| ΔCC (h*IMMT*<sup>432-697</sup>) | This paper | |
| ΔMD (h*IMMT*<sup>698-758</sup>) | This paper | |
| K299E (h*IMMT*<sup>A895G</sup>) | This paper |  |
| Software and algorithms |  |  |
| ImageStudio, v6.1 | Li-COR |  |
| FIJI v1.54p | NIH |  |
| Huygens Pro, v25.10 | Scientific Volume Imaging |  |
| LASX, v4.8.1.29271 | Leica |  |
| Zen Black, v3.0 | Zeiss |  |
| Imaris, v10.2.0 | Imaris |  |
| ilastik, v1.4.1rc2 | Ilastik |  |
| Wave, v2.6.4 | Agilent Technologies |  |
| Cell Imaging, v1.2.0.12 | Agilent Technologies |  |
| PyMOL, v2.4.0 | Schrodinger |  |
| ViCell Blu Cell Viability Analyzer, v1.4.4.1 | Beckman Coulter |  |
| MitoMeter | Lefebvre *et al.*<sup>44</sup> |  |
| Prism, v10.5.0 | GraphPad |  |
| R, v4.4.2 | R Core Team |  |
| https://github.com/stjude-cbi/mito-analysis | This paper |  |
| Other |  |  |

### Experimental model and study participant details

#### Mice

We utilized our previously described conditional mouse model targeting *Immt*^16^ for this study. The number of mice used for each experiment is detailed in the respective figure legends. Both male and female mice were used; no sex-dependent differences in deletion efficiency were identified in our previous characterization of this model^16^, and data from both sexes were pooled for all analyses. AAV containing a liver protein 1 (LP1) promoter to drive expression of Cre recombinase (LP1-Cre AAV) was generated by the Vector Core Laboratory at SJCRH (Memphis, TN, USA). As previously described^27^, stock AAV was diluted to 2×10^11^ gc/100 µL in sterile phosphate buffered saline (PBS), and 100 µL was injected into the tail vein of recipient mice (ages ranged from 9 to 32 weeks at the time of injection). Mice were monitored for health, and serum from retro-orbital blood was collected every week for chemical analysis. Serum levels for ALT and AST were assessed with the Pentra C400 Clinical Chemistry Analyzer per the manufacturer’s instructions (HORIBA, Irvine, CA, USA). Mice were housed with food and water provided *ad libitum* on a 12 h light/dark cycle. A visual assessment of hunched appearance, scruffy fur, and minimal movement determined the humane endpoint^58^. All procedures were approved by the St. Jude Institutional Animal Care and Use Committee (Protocol 2384).

#### Cell culture

MEFs were prepared from *Immt*^WT^, *Immt*^f/WT^, or *Immt*^f/f^ embryos (taken at E12–13)^59^ and immortalized with SV40 linearized plasmid. HEK293T cells were obtained from the ATCC (CRL-3216) and used solely for virus production. All cells were maintained in Dulbecco’s Modified Eagle Medium (DMEM; high glucose) supplemented with 8% fetal bovine serum (FBS), 1% penicillin/streptomycin, 1% L-glutamine, and 1% non-essential amino acids. To delete the floxed *Immt* allele in MEFs, cells were treated with 100 nM 4-hydroxy-tamoxifen (4-OH-TAM, H7904 Sigma Aldrich) or equivalent DMSO. The day before harvesting cells for analyses, MEFs were re-seeded at ∼5000 cells/cm^2^ in respective treatment media. Cells were assessed for quantity and viability using the Vi-Cell Blu Cell Viability Analyzer (Beckman Coulter, Brea, CA, USA). All cell lines were maintained at 37°C with 5% CO_2_.and verified to be mycoplasma negative throughout the duration of these studies.

For lentivirus production, HEK293T cells were seeded at 65,000 cells/cm^2^, then transfected with 12 µg plasmid DNA using Fugene-6 (Promega, Madison, WI, USA). Lentiviral particles were collected between 36–72 h post-transfection, filtered (0.45 µm), and applied to target MEFs (*Immt*^f/f^ ROSA-CreER^T2^) for infection. After at least one round of passaging, the top 20% of zsGreen-positive MEFs were sorted (Flow Cytometry Core, SJCRH, Memphis, TN, USA) and used for downstream analyses.

For retrovirus production, HEK293T cells were seeded at 65,000 cells/cm^2^, then transfected with 12 µg plasmid DNA (pBabe-puro-IMS-RP, a gift from Peter Sorger (Addgene plasmid # 24535)^36^ and retroviral packaging plasmids) using Fugene-6 (Promega, Madison, WI, USA). Retroviral particles were collected between 36–72 h post-transfection, filtered (0.45 µm), and applied to target MEFs (*Immt*^f/f^ ROSA-CreER^T2^) for infection. Forty-eight hours after infection, MEFs were positively selected with 1 µg/ml puromycin.

### Method details

#### Immunohistochemical studies

When LP1-Cre AAV–injected mice reached an endpoint, they were euthanized via CO_2_. For immunohistochemistry, liver tissue samples were excised, rinsed in PBS, and fixed in 4% neutral buffered formalin. All samples were provided to the Comparative Pathology Core (SJCRH, Memphis, TN, USA) for paraffin embedding, slicing, and staining with hematoxylin & eosin or the indicated antibody (see the STAR Methods for antibody details). For all experiments, slides were independently analyzed by a certified pathologist as previously described^16^.

#### Cloning

We first obtained lentiviral particles containing the cDNA for h*IMMT* from Origene. WT MEFs (lacking ROSA-CreER^T2^) were infected with the lentivirus, followed by genomic DNA isolation and PCR to amplify the human cDNA. h*IMMT* was then subcloned into pCR-II TOPO vector using primers to add a V5 tag to the C-terminus and NotI and BamHI restriction enzyme sites (all primer sequences are described in **Supplementary Table 5**). After Sanger sequencing validation for sequence fidelity, h*IMMT* was transferred into the pHAGE-UBC-MCS-IRES-ZsGreen-W lentiviral backbone to provide a fluorescent reporter for infected cells.

Primers were designed to excise targeted regions of h*IMMT* using overlap extension PCR as follows: ΔTM (transmembrane domain, aa 41–62), ΔHB (helical bundle, aa 203–432), ΔCC (coiled-coil, aa 433–697), and ΔMD (mitofilin domain, aa 698–758). All primer sequences are detailed in **Supplementary Table 5**. The first round of PCR was completed using 500-ng full-length h*IMMT* in pCR-II TOPO, 200 μM dNTPs, 1x Q5 Reaction Buffer, 1x Q5 GC Enhancer, 0.1 μM each of forward and reverse primers, and Q5 polymerase. The PCR reaction ran as follows: 1 cycle of 94°C for 1 min, 20 cycles of a) 94°C for 30 s, b) 55°C for 30 s, and c) 72°C for 150 s, and finally, 1 cycle of 72°C for 10 min. PCR products from the first round, containing at least a 30-nt overlapping sequence, were gel purified and combined at a 1:1 (DNA mass) ratio for the second round of PCR with the h*IMMT* forward and V5 reverse primers. The second PCR reaction ran as follows: 1 cycle of 94°C for 5 min, 38 cycles of a) 94°C for 30 s, b) 55°C for 30 s, and c) 72°C for 150 s, and finally 1 cycle of 72°C for 10 min. The resulting PCR product was gel purified and ligated to Zero Blunt pCR™-Blunt II-TOPO (Invitrogen) per the manufacturer’s protocol. After sequence validation for the respective mutations, the cDNA was excised from TOPO using NotI and BamHI restriction enzymes, then ligated to pHAGE-UBC-MCS-IRES-ZsGreen-W as detailed above. Sequence fidelity of the constructs was validated using Sanger sequencing.

Site-directed mutagenesis to generate the K299E mutant was completed using the QuikChange II XL Site-Directed Mutagenesis Kit (Agilent). Primers were designed according to Agilent’s QuikChange Primer Design Tool (see **Supplemental Table 5**) to create the 895A>G point mutation. Full-length h*IMMT* with the V5 tag in pCRII-TOPO was used as the template, and PCR reaction ran as follows: 1 cycle of 95°C for 1 min, 18 cycles of a) 95°C for 50 s, b) 60°C for 50 s, and c) 68°C for 10 min, and finally 1 cycle of 68°C for 7 min. The reaction was digested with Dpn1 to remove the template DNA before transforming XL10 Gold cells. After validating the mutation, the K299E cDNA was moved to the pHAGE-UBC-MCS-IRES-ZsGreen-W lentiviral backbone as detailed above.

#### Immunoblotting

Protein levels from tissues and cells were assessed as previously described^16, 60^. To assess the OPA1 isoforms, samples were run on a 3%–8% tris-acetate SDS-PAGE gel in cold tris-acetate running buffer, then transferred to polyvinylidene fluoride membranes at 20 V for 40 min. Membranes were cut horizontally to assess proteins of varying molecular weights and incubated overnight at 4°C in primary antibodies diluted in 5% BSA-PBST (BSA, BP1600, Thermo Fisher Scientific). The STAR Methods table summarizes all primary and secondary antibodies used for western blotting analyses. Chemiluminescence was read on a Li-COR Odyssey Fc Imager (Li-COR Biosciences, Lincoln, NE, USA) using ImageStudio (v6.1). No changes were made to image brightness or contrast after acquisition, and if required, blots were adjusted for horizontal alignment using FIJI (version 1.54p, NIH, Bethesda, MD, USA).

#### IP assay

Approximately 20×10^6^ cells were harvested as previously described^61^. Then, 1.5 mg total protein lysate was incubated with 3 μg of V5 antibody for 4 h at 4°C before Protein A/G Magnetic Beads (Invitrogen, Waltham, MA, USA) were added to incubate overnight at 4°C. The next day, beads were washed with cold 1x PBS and protein was eluted with 1x LDS Sample Buffer + 10% dithiothreitol with a 5 min boil at 95°C.

#### Mitochondrial isolation

Mitochondria were isolated with the Miltenyi Mitochondrial Isolation Kit per the manufacturer’s instructions. Briefly, 50–100 mg of minced liver tissue or 20–30 ×10^6^ cells were placed in a Dounce and homogenized with 2 mL ice-cold Lysis Buffer for 10–15 strokes. Homogenized samples were next passed through a 30-µm Pre-Separation Filter (120-009-491, Miltenyi). The 2-mL filtered sample was diluted to 10 mL with cold 1x Separation Buffer; then, 50 µL anti-TOM22 magnetic MicroBeads were added to each sample to incubate for at least 1 h on a rotator at 4°C. Samples were passed through LS Columns (130-042-401, Miltenyi) placed inside the QuadroMACS Separation Unit (130-091-051, Miltenyi) to isolate the magnetically tagged mitochondria. Samples were eluted with 1.5 mL 1x Separation Buffer, then pelleted at 5000 *g* for 5 min (4°C) and suspended in an appropriate volume of ice cold 1x Separation Buffer per the instructions. Mitochondrial concentration was assessed using the BCA Protein Assay Kit.

#### MOMP assay

The MOMP assay was completed on isolated mitochondria as previously described^62, 63^. Briefly, 50 µg of mitochondria was suspended in mitochondrial assay buffer, then incubated for 1 h at 37°C while increasing concentrations of the BID-stapled alpha-helical peptide SAHB^30^. Samples were centrifuged at 5500 *g* for 5 min, the supernatant was collected in 2x sample buffer, and the pellet was solubilized in 1x sample buffer. Samples ran on SDS-PAGE to assess cytochrome *c*.

#### MICOS assembly assay (2D blue native/SDS-PAGE)

The MICOS assembly assay was completed as previously described^38^, with modifications to assess mitochondrial supercomplexes^64^. The BCA Protein Assay Kit was used to determine protein concentration from isolated mitochondria, the pellets of which were lysed in 4 g/g digitonin prepared in native-PAGE sample buffer (Invitrogen, Waltham, MA, USA). The crude fraction was centrifuged out, and G-250 sample additive was added to each sample. A sample containing 100 µg of protein was loaded to each lane of a 3%–12% NativePAGE gel (Invitrogen, Waltham, MA, USA). Samples were run in Dark Blue Cathode Running Buffer (prepared with 1:20 Cathode Buffer Additive, BN2002, Thermo Fisher Scientific) for 20 min, then in Light Blue Cathode Running Buffer (prepared with 1:200 Cathode Buffer Additive) until the dye front reached the bottom of the gel. Individual lanes were excised and incubated in denaturing buffer^38^, then placed in the well of a 10% NuPAGE 2D-well gel. Western blots were processed as described above.

#### Agilent Seahorse assays

*Immt* ROSA-CreER^T2^ MEFs were treated with DMSO or 100 nM TAM. The day before analysis, MEFs were seeded at 25,000 cells/well in the XF 96 Cell Culture Microplate (Agilent, Santa Clara, CA, USA) in the respective treatment. The next morning, Seahorse media was freshly prepared per manufacturer’s instructions (Seahorse DMEM Base Media without Phenol Red, pH 7.4 (103575), 1% L-Glutamine, 1% Sodium Pyruvate, 10 mM D-Glucose) and warmed to 37°C. The media from the plate was gently discarded before the warmed Seahorse media was carefully added to each well to avoid detaching cells from the base. The cells were first imaged on a BioTek Cytation5 Imaging Reader (v1.2.0.12, BioTek, Winooski, VT, USA), and the cartridge ports were prepared with 1.5 µM oligomycin, 8 µM carbonyl cyanide-p-trifluoromethoxyphenylhydrazone (FCCP), and 0.5 µM rotenone/Antimycin A plus Hoechst dye (all prepared in Seahorse media). Oxygen consumption rate was assessed using the Agilent MitoStress protocol on the Wave software (v2.6.4). At the end of the assay, the cell plate was read on the BioTek Cytation5 for fluorescence to normalize the results by cell counts per well.

#### Fixed cell super-resolution SIM

*Immt* ROSA-CreER^T2^ MEFs were treated with DMSO or 100 nM TAM. The day before analysis, MEFs were seeded at 5,000 cells/cm^2^ on No. 1.5H 10-mm cover glass (64-0718, Warner Instruments, Hamden, CT, USA) in complete media with the respective treatment. The next day, cells were rinsed with warm (37°C) 1x PHEM Buffer (60 mM PIPES, 25 mM HEPES, 10 mM EGTA, and 4 mM MgCl^2^, pH 6.9) before fixing for 15 min in 4% paraformaldehyde-PHEM (37°C). After a rinse with 1x PHEM, cells were gently permeabilized with 0.1% Triton X-100-PHEM for 10 min. The samples were rinsed again with 1x PHEM, then blocked at 25°C in 5% FBS-PHEM for 1 h. After a final wash with 1x PHEM, the primary antibody cocktail (containing 1:250 TOM20 and cytochrome *c* antibodies, 1% FBS, and 1x PHEM) was added to the wells and the samples incubated at 4°C overnight. The next day, the samples were rinsed with 1x PHEM, then secondary antibody cocktail containing the respective AlexaFluor fluorescent antibodies (1:250), 1% FBS, and 1x PHEM was added to samples before incubation at 4°C overnight. On the last day, cells were washed three times with 1x PHEM (the last wash containing 1:500 DAPI) before mounting to glass slides with Prolong Diamond Antifade Mountant (P36965, Invitrogen). Samples cured for at least 48 h before imaging on an Elyra7 (Zeiss, Oberkochen, Germany) for Lattice SIM imaging (63× 1.4 NA objective). SIM^2^ post-processing was performed in ZEN Black (v3.0, Zeiss) using weak SNR settings and each channel was aligned and registered using a 100nm TetraSpeck Microspheres (Thermo Fisher Scientific) acquired and processed using similar settings.

#### Live-cell STED microscopy

For live-cell imaging, *Immt* ROSA-CreER^T2^ MEFs were treated with DMSO or 100 nM TAM. The day before analysis, MEFs were seeded at 5,000 cells/cm^2^ on ibidi No. 1.5H glass bottom dishes or 8-well chamber slides (81158 and 80807, respectively, ibidi, Gräfelfing, Germany) and then returned to 37°C to adhere overnight. The next day, MEFs were treated with 1:1000 Spirochrome PK-Mito ORANGE Dye (CY-SC053 (Cytoskeleton, Inc, Denver, CO, USA) and 1:1000 Quant-iT PicoGreen dsDNA dye (P11495, Invitrogen), prepared in imaging media (IM; DMEM without Phenol Red, 8% FBS, 1% penicillin-streptomycin, 1% L-Glutamine, 1% MEM-NEAA) for 1 h at 37°C. Cells were rinsed with IM, and IM was maintained through the duration of the imaging experiment. Live cells were housed on the STELLARIS 8 tauSTED (Leica, Wetzler, Germany) for imaging with the 100× 1.4 NA objective at 37°C and 5% CO_2_. Single z-plane images were captured at 28 nm/pixel with a scanner frequency of 400 Hz, 16 lines accumulation and simultaneous channel acquisition with STED depletion at 775nm using previously published excitation/emission wavelengths for PKMO^33^ and picoGreen^65^.

#### Live-cell confocal microscopy

To assess the inner contents of the rounded mitochondria, IMS-RP–expressing *Immt* ROSA-CreER^T2^ MEFs were treated with 100 nM TAM. The day before analysis, MEFs were seeded at 5,000 cells/cm^2^ on ibidi No. 1.5H glass bottom 8-well chamber slides. The next day, MEFs were treated with 1:1000 Spirochrome PK-Mito DEEP RED Dye (PKMDR, CY-SC055 (Cytoskeleton, Inc, Denver, CO, USA) and 1:1000 Quant-iT PicoGreen dsDNA dye (P11495, Invitrogen), for 1 h at 37°C. Cells were rinsed with IM, which was maintained through the duration of the imaging experiment. Live cells were housed on the STELLARIS 8 (Leica, Wetzler, Germany) for imaging with the 100× 1.4 NA lens at 37°C and 5% CO_2_. Z-stacks were captured at every 260 nm along the Z-axis, with a pixel size of 30 nm/pixel with a scanner frequency of 100 Hz, and 2 line accumulation. The 494 nm (PicoGreen) and 649 nm (PKMDR) channels were acquired simultaneously while the 584 nm (IMS-RP) channel was acquired sequentially.

To assess mitochondrial membrane potential, *Immt* ROSA-CreER^T2^ MEFs were treated with 100 nM TAM. The day before analysis, MEFs were seeded at 5,000 cells/cm^2^ on ibidi No. 1.5H glass bottom 8-well chamber slides then returned to 37°C to adhere overnight. The next day, MEFs were treated with 200 nM TMRM and 500 nM MitoTracker Green (M7514, Invitrogen) before incubation at 37°C for 30 min. Cells were then rinsed with IM, which was maintained for the duration of the imaging experiment. Live cells were housed on the STELLARIS 8 (Leica, Wetzler, Germany) for imaging with the 100× 1.4 NA at 37°C and 5% CO_2_. Single z-plane images were captured at 28 nm/pixel with a scanner frequency at a pixel size of 60 nm/pixel with a scanner frequency of 400 Hz, 16 lines averaging and simultaneous channel acquisition.

#### Mitochondrial quantification pipeline

TauSTED images of PKMO-stained MEFs were deconvolved using a theoretical STED point spread function for STED in Huygens Pro (v25.10, Scientific Volume Imaging, Hilversum, The Netherlands). Mitochondrial objects were then segmented using a 2D U-Net model trained within the nnU-Net framework^66^. A total of 21 fields of view (FOVs) were manually annotated to generate ground-truth binary masks for model training. Following segmentation, a boundary-to-boundary skeletonization procedure was applied to trace midlines through each mitochondrial object, in which skeleton endpoints were extended to the nearest object boundary. The resulting skeleton mask was then applied to the original deconvolved image, and the NeuroCyto tool^67^ was used to extract intensity measurements along each line. The developed quantification pipeline is available at: https://github.com/stjude-cbi/mito-analysis.

#### STEM and FIBSEM

For scanning electron transmission microscopy (STEM), LP1-Cre treated mice were perfused with PBS followed by electron microscopy-grade 4% paraformaldehyde in 0.1 M phosphate buffer (pH 7.2). Liver sections were excised and flushed with EM fixative, then processed and analyzed as previously described^16^.

For FIBSEM, mitochondria were isolated from liver tissues as detailed above, then pelleted and fixed with EM fixative (2.5% glutaraldehyde, 2% PFA in 0.1 M phosphate buffer). Pelleted samples were enrobed in 4% low melting point agarose and processed as previously described^16^. Samples were examined in a Thermo Fisher Scientific Helios Nanolab G3 FIBSEM. Image stacks were aligned, and Ilastik (v1.4.1rc2) was used for mitochondrial segmentation. The FIBSEM stack and prediction outputs from Ilastik were loaded to Imaris (v10.2.0) to clean up the segmentation, analyze the mitochondrial volume, and generate the final images and videos.

### Quantification and statistical analyses

To evaluate treatment-level differences in mitochondrial features, we analyzed segmented mitochondria from microscopy images using linear mixed models (LMM) that accounted for the hierarchical data structure, with multiple mitochondria per image and multiple images per biological replicate nested within treatment. Random intercepts were included for image and biological replicate to capture clustering.

Prior to analysis, feature values were winsorized at the 99.5th percentile to reduce the influence of extreme observations. Most features (except solidity, gray level co-occurrence matrix (glcm) homogeneity mean, and glcm correlation mean) were log-transformed to better approximate normality and stabilize variance. Distributional assumptions were assessed, and the analyses were considered robust to minor deviations from normality. Mitochondrial features were grouped into three categories: morphology (solidity, circularity, area-to-perimeter ratio, major axis length, minor axis length), intensity (mean intensity, max intensity), and texture (glcm contrast mean, glcm homogeneity mean, glcm correlation mean). For each feature, treatment effects were estimated using LMM of the form:

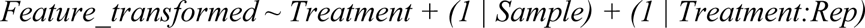

where *Sample* and *Rep* represent the image and replicate identifiers, respectively, with replicates nested within treatment. This model accounts for correlations among mitochondria from the same image and among images from the same replicate.

Two sets of treatment comparisons were performed: (i) *Immt*^f/f^ versus the wild-type (WT) reference and (ii) full-length h*IMMT*, ΔTM, ΔHB, ΔCC, ΔMD, and K299E versus the empty vector (EV) control. Model coefficients represent estimated treatment effects relative to the corresponding reference group. For log-transformed features, estimates were back-transformed and reported as percentage changes. Multiple testing was controlled using the Benjamini–Hochberg false discovery rate (FDR), applied within each feature category across treatment comparisons. All tests were two-sided with a significance level of 0.05. Effect estimates and 95% confidence intervals were obtained from the fitted models. Analyses were conducted in R (Version 4.4.2, R Foundation for Statistical Computing), and results were visualized using effect estimates with confidence intervals for each feature and treatment comparisons.

All graphs were prepared in GraphPad Prism (Version 9.5.1(528), San Diego, CA, USA). For murine survival studies, a Log-rank Mantel-Cox test, followed by a Gehan, Breslow, Wilcoxon test was completed. For statistical analyses of the parental (*Immt*^WT^ and *Immt*^Δ/Δ^) MEFs, an unpaired Welch’s *t*-test was used. For statistical analyses of the h*IMMT* mutants, a one-way multi-comparison ANOVA was completed for comparison to the EV MEFs. For all graphs, results are presented as the mean ± standard error (SEM). For all fluorescent microscopy experiments, the sample size calculator (https://homepage.univie.ac.at/robin.ristl/samplesize.php?test=ttest) was used to determine appropriate sample size (two-sided alpha = 0.05, power = 0.8) using comparisons between WT and *Immt*^Δ/Δ^ (parental MEFs, **Figure 2**) or EV and h*IMMT* (mutant MEFs, **Figure 4**) MEFs. Statistical significance was set to: * *p* ≤0.05, ** *p* ≤ 0.01, *** *p* ≤0.001, **** *p* ≤ 0.0001, ns = not significant.

## References

1. McBride, H.M., M. Neuspiel, and S. Wasiak, Mitochondria: more than just a powerhouse. Curr Biol, 2006. 16(14): p. R551–60.

2. Davies, K.M., et al., Structure of the yeast F1Fo-ATP synthase dimer and its role in shaping the mitochondrial cristae. Proc Natl Acad Sci U S A, 2012. 109(34): p. 13602–7.

3. Glancy, B., et al., The Functional Impact of Mitochondrial Structure Across Subcellular Scales. Front Physiol, 2020. 11: p. 541040.

4. Friedman, J.R. and J. Nunnari, Mitochondrial form and function. Nature, 2014. 505(7483): p. 335–43.

5. Eramo, M.J., et al., The ‘mitochondrial contact site and cristae organising system’ (MICOS) in health and human disease. J Biochem, 2020. 167(3): p. 243–255.

6. Diederichs, K.A., et al., Structural insight into mitochondrial beta-barrel outer membrane protein biogenesis. Nat Commun, 2020. 11(1): p. 3290.

7. Huynen, M.A., et al., Evolution and structural organization of the mitochondrial contact site (MICOS) complex and the mitochondrial intermembrane space bridging (MIB) complex. Biochim Biophys Acta, 2016. 1863(1): p. 91–101.

8. John, G.B., et al., The mitochondrial inner membrane protein mitofilin controls cristae morphology. Mol Biol Cell, 2005. 16(3): p. 1543–54.

9. Rabl, R., et al., Formation of cristae and crista junctions in mitochondria depends on antagonism between Fcj1 and Su e/g. J Cell Biol, 2009. 185(6): p. 1047–63.

10. Zerbes, R.M., et al., Role of MINOS in mitochondrial membrane architecture: cristae morphology and outer membrane interactions differentially depend on mitofilin domains. J Mol Biol, 2012. 422(2): p. 183–91.

11. Ott, C., et al., Detailed analysis of the human mitochondrial contact site complex indicate a hierarchy of subunits. PLoS One, 2015. 10(3): p. e0120213.

12. Li, H., et al., Mic60/Mitofilin determines MICOS assembly essential for mitochondrial dynamics and mtDNA nucleoid organization. Cell Death Differ, 2016. 23(3): p. 380–92.

13. Kondadi, A.K., et al., Cristae undergo continuous cycles of membrane remodelling in a MICOS-dependent manner. EMBO Rep, 2020. 21(3): p. e49776.

14. Feng, Y., et al., Mitofilin Heterozygote Mice Display an Increase in Myocardial Injury and Inflammation after Ischemia/Reperfusion. Antioxidants (Basel), 2023. 12(4).

15. Dong, T., et al., Mic60 is essential to maintain mitochondrial integrity and to prevent encephalomyopathy. Brain Pathol, 2023. 33(4): p. e13157.

16. Rockfield, S.M., et al., Genetic ablation of Immt induces a lethal disruption of the MICOS complex. Life Sci Alliance, 2024. 7(6).

17. Bock-Bierbaum, T., et al., Structural insights into crista junction formation by the Mic60-Mic19 complex. Sci Adv, 2022. 8(35): p. eabo4946.

18. Odgren, P.R., et al., Molecular characterization of mitofilin (HMP), a mitochondria-associated protein with predicted coiled coil and intermembrane space targeting domains. J Cell Sci, 1996. 109 (Pt 9): p. 2253–64.

19. Gieffers, C., et al., Mitofilin is a transmembrane protein of the inner mitochondrial membrane expressed as two isoforms. Exp Cell Res, 1997. 232(2): p. 395–9.

20. Korner, C., et al., The C-terminal domain of Fcj1 is required for formation of crista junctions and interacts with the TOB/SAM complex in mitochondria. Mol Biol Cell, 2012. 23(11): p. 2143–55.

21. Zhu, C., et al., Single-molecule, full-length transcript isoform sequencing reveals disease-associated RNA isoforms in cardiomyocytes. Nat Commun, 2021. 12(1): p. 4203.

22. Daumke, O. and M. van der Laan, Molecular machineries shaping the mitochondrial inner membrane. Nat Rev Mol Cell Biol, 2025. 26(9): p. 706–724.

23. Marco-Hernandez, A.V., et al., Mitochondrial developmental encephalopathy with bilateral optic neuropathy related to homozygous variants in IMMT gene. Clin Genet, 2022. 101(2): p. 233–241.

24. Tirrell, P.S., et al., MICOS subcomplexes assemble independently on the mitochondrial inner membrane in proximity to ER contact sites. J Cell Biol, 2020. 219(11).

25. Hessenberger, M., et al., Regulated membrane remodeling by Mic60 controls formation of mitochondrial crista junctions. Nat Commun, 2017. 8: p. 15258.

26. Tsai, P.I., et al., PINK1 Phosphorylates MIC60/Mitofilin to Control Structural Plasticity of Mitochondrial Crista Junctions. Mol Cell, 2018. 69(5): p. 744–756 e6.

27. Wright, T., et al., Anti-apoptotic MCL-1 promotes long-chain fatty acid oxidation through interaction with ACSL1. Mol Cell, 2024. 84(7): p. 1338–1353 e8.

28. Nallagangula, K.S., et al., Liver fibrosis: a compilation on the biomarkers status and their significance during disease progression. Future Sci OA, 2018. 4(1): p. FSO250.

29. Yang, R.F., et al., Mitofilin regulates cytochrome c release during apoptosis by controlling mitochondrial cristae remodeling. Biochem Biophys Res Commun, 2012. 428(1): p. 93–8.

30. Zheng, J.H., et al., Intrinsic Instability of BOK Enables Membrane Permeabilization in Apoptosis. Cell Rep, 2018. 23(7): p. 2083–2094 e6.

31. Madungwe, N.B., et al., Mitochondrial inner membrane protein (mitofilin) knockdown induces cell death by apoptosis via an AIF-PARP-dependent mechanism and cell cycle arrest. Am J Physiol Cell Physiol, 2018. 315(1): p. C28–C43.

32. Gu, X., et al., Measurement of mitochondrial respiration in adherent cells by Seahorse XF96 Cell Mito Stress Test. STAR Protoc, 2021. 2(1): p. 100245.

33. Liu, T., et al., Multi-color live-cell STED nanoscopy of mitochondria with a gentle inner membrane stain. Proc Natl Acad Sci U S A, 2022. 119(52): p. e2215799119.

34. Ashley, N., D. Harris, and J. Poulton, Detection of mitochondrial DNA depletion in living human cells using PicoGreen staining. Exp Cell Res, 2005. 303(2): p. 432–46.

35. Povea-Cabello, S., M. Brischigliaro, and E. Fernandez-Vizarra, Emerging mechanisms in the redox regulation of mitochondrial cytochrome c oxidase assembly and function. Biochem Soc Trans, 2024. 52(2): p. 873–885.

36. Albeck, J.G., et al., Modeling a snap-action, variable-delay switch controlling extrinsic cell death. PLoS Biol, 2008. 6(12): p. 2831–52.

37. Yang, Z., et al., Cyclooctatetraene-conjugated cyanine mitochondrial probes minimize phototoxicity in fluorescence and nanoscopic imaging. Chem Sci, 2020. 11(32): p. 8506–8516.

38. Kumar, A., et al., A dynamin superfamily-like pseudoenzyme coordinates with MICOS to promote cristae architecture. Curr Biol, 2024. 34(12): p. 2606–2622 e9.

39. Barrera, M., et al., OPA1 functionally interacts with MIC60 but is dispensable for crista junction formation. FEBS Lett, 2016. 590(19): p. 3309–3322.

40. Darshi, M., et al., ChChd3, an inner mitochondrial membrane protein, is essential for maintaining crista integrity and mitochondrial function. J Biol Chem, 2011. 286(4): p. 2918–32.

41. Glytsou, C., et al., Optic Atrophy 1 Is Epistatic to the Core MICOS Component MIC60 in Mitochondrial Cristae Shape Control. Cell Rep, 2016. 17(11): p. 3024–3034.

42. Elachouri, G., et al., OPA1 links human mitochondrial genome maintenance to mtDNA replication and distribution. Genome Res, 2011. 21(1): p. 12–20.

43. Macuada, J., et al., OPA1 and disease-causing mutants perturb mitochondrial nucleoid distribution. Cell Death Dis, 2024. 15(11): p. 870.

44. Lefebvre, A., et al., Automated segmentation and tracking of mitochondria in live-cell time-lapse images. Nat Methods, 2021. 18(9): p. 1091–1102.

45. Nathanail, E., et al., Integrative modelling reveals the structure of the human Mic60-Mic19 subcomplex and its role as a diffusion barrier in mitochondria. bioRxiv, 2026: p. 2026.01.30.702776.

46. Deng, R., et al., Loss of MIC60 Aggravates Neuronal Death by Inducing Mitochondrial Dysfunction in a Rat Model of Intracerebral Hemorrhage. Mol Neurobiol, 2021. 58(10): p. 4999–5013.

47. Ikeda, H., et al., Miclxin, a Novel MIC60 Inhibitor, Induces Apoptosis via Mitochondrial Stress in beta-Catenin Mutant Tumor Cells. ACS Chem Biol, 2020. 15(8): p. 2195–2204.

48. Ge, Y., et al., Two forms of Opa1 cooperate to complete fusion of the mitochondrial inner-membrane. Elife, 2020. 9.

49. Del Dotto, V., et al., Eight human OPA1 isoforms, long and short: What are they for? Biochim Biophys Acta Bioenerg, 2018. 1859(4): p. 263–269.

50. Zeharia, A., et al., Mitochondrial hepato-encephalopathy due to deficiency of QIL1/MIC13 (C19orf70), a MICOS complex subunit. Eur J Hum Genet, 2016. 24(12): p. 1778–1782.

51. Godiker, J., et al., QIL1-dependent assembly of MICOS complex-lethal mutation in C19ORF70 resulting in liver disease and severe neurological retardation. J Hum Genet, 2018. 63(6): p. 707–716.

52. Kishita, Y., et al., A novel homozygous variant in MICOS13/QIL1 causes hepato-encephalopathy with mitochondrial DNA depletion syndrome. Mol Genet Genomic Med, 2020. 8(10): p. e1427.

53. Wang, Z.A. and P.A. Cole, The Chemical Biology of Reversible Lysine Post-translational Modifications. Cell Chem Biol, 2020. 27(8): p. 953–969.

54. Akabane, S., et al., PKA Regulates PINK1 Stability and Parkin Recruitment to Damaged Mitochondria through Phosphorylation of MIC60. Mol Cell, 2016. 62(3): p. 371–384.

55. Bomba-Warczak, E., et al., Long-lived mitochondrial cristae proteins in mouse heart and brain. J Cell Biol, 2021. 220(9).

56. Krishna, S., et al., Identification of long-lived proteins in the mitochondria reveals increased stability of the electron transport chain. Dev Cell, 2021. 56(21): p. 2952–2965 e9.

57. Mootha, V.K., et al., Integrated analysis of protein composition, tissue diversity, and gene regulation in mouse mitochondria. Cell, 2003. 115(5): p. 629–40.

58. Burkholder, T., et al., Health Evaluation of Experimental Laboratory Mice. Curr Protoc Mouse Biol, 2012. 2: p. 145–165.

59. Conner, D.A., Mouse embryo fibroblast (MEF) feeder cell preparation. Curr Protoc Mol Biol, 2001. Chapter 23: p. Unit 23 2.

60. Stewart, D.P., et al., Ubiquitin-independent degradation of antiapoptotic MCL-1. Mol Cell Biol, 2010. 30(12): p. 3099–110.

61. Stephan, T., et al., MICOS assembly controls mitochondrial inner membrane remodeling and crista junction redistribution to mediate cristae formation. EMBO J, 2020. 39(14): p. e104105.

62. Renault, T.T., K.V. Floros, and J.E. Chipuk, BAK/BAX activation and cytochrome c release assays using isolated mitochondria. Methods, 2013. 61(2): p. 146–55.

63. Singh, G. and T. Moldoveanu, Methods to Probe Conformational Activation and Mitochondrial Activity of Proapoptotic BAK. Methods Mol Biol, 2019. 1877: p. 185–200.

64. Jha, P., X. Wang, and J. Auwerx, Analysis of Mitochondrial Respiratory Chain Supercomplexes Using Blue Native Polyacrylamide Gel Electrophoresis (BN-PAGE). Curr Protoc Mouse Biol, 2016. 6(1): p. 1–14.

65. Stephan, T., et al., Live-cell STED nanoscopy of mitochondrial cristae. Sci Rep, 2019. 9(1): p. 12419.

66. Isensee, F., et al., nnU-Net: a self-configuring method for deep learning-based biomedical image segmentation. Nat Methods, 2021. 18(2): p. 203–211.

67. Culley, S., et al., Quantitative mapping and minimization of super-resolution optical imaging artifacts. Nat Methods, 2018. 15(4): p. 263–266.

