## Supplemental Tables, Figures, Legends for "Protein domain characterization reveals human MIC60 tolerates loss of helical bundle domain"

### Supplemental Figure 1

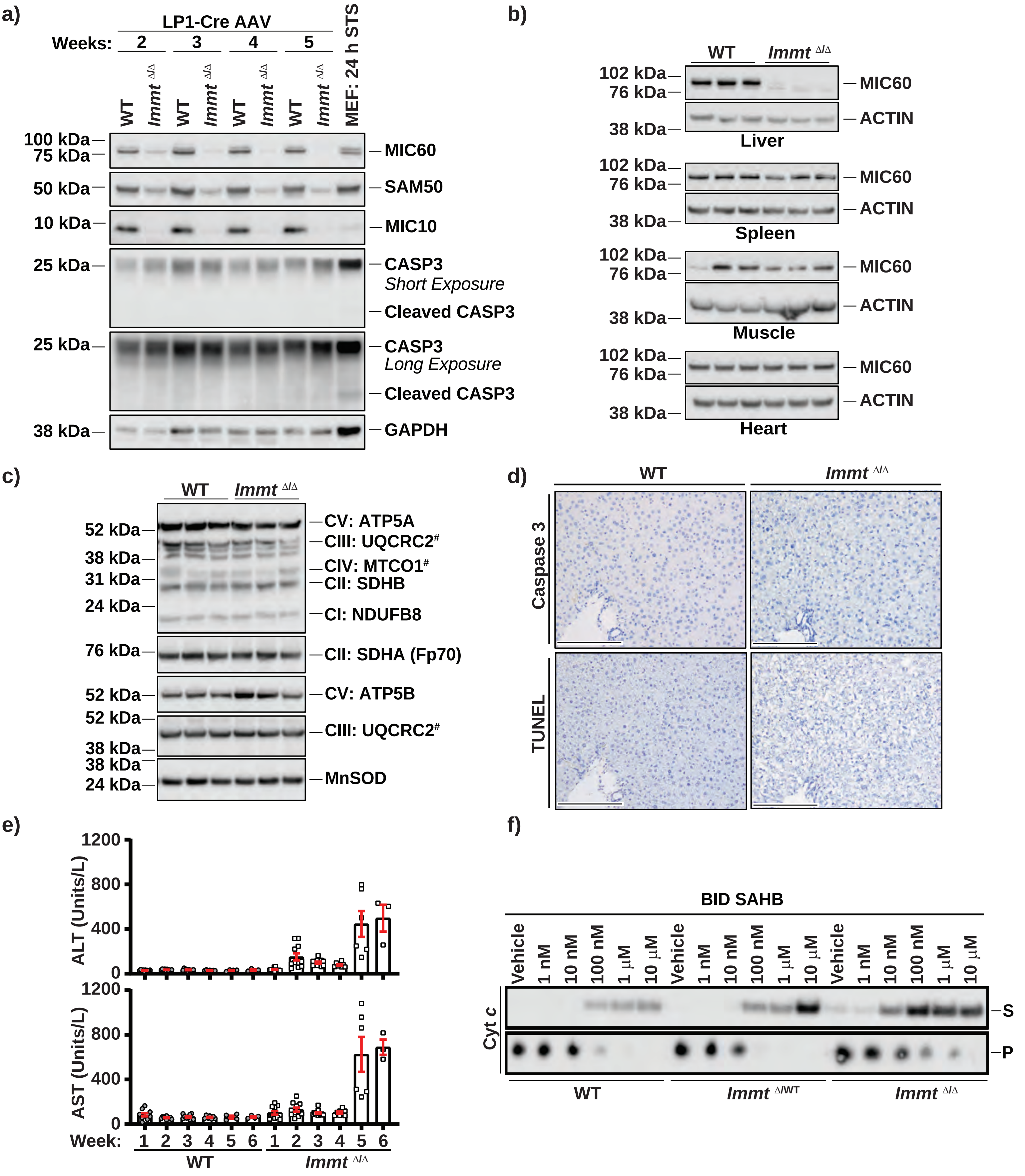

#### Supplemental Figure 2

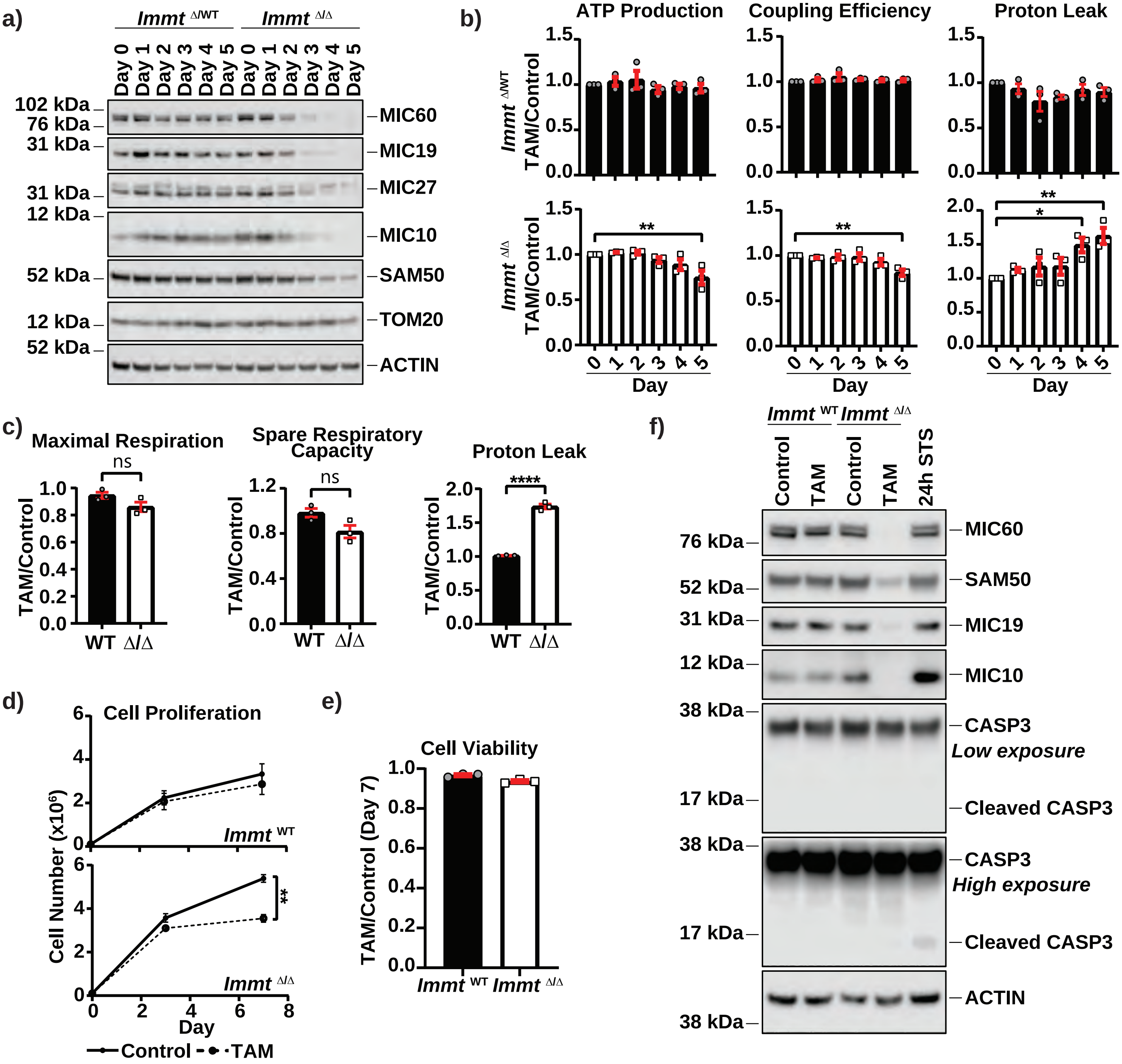

#### Supplemental Figure 3

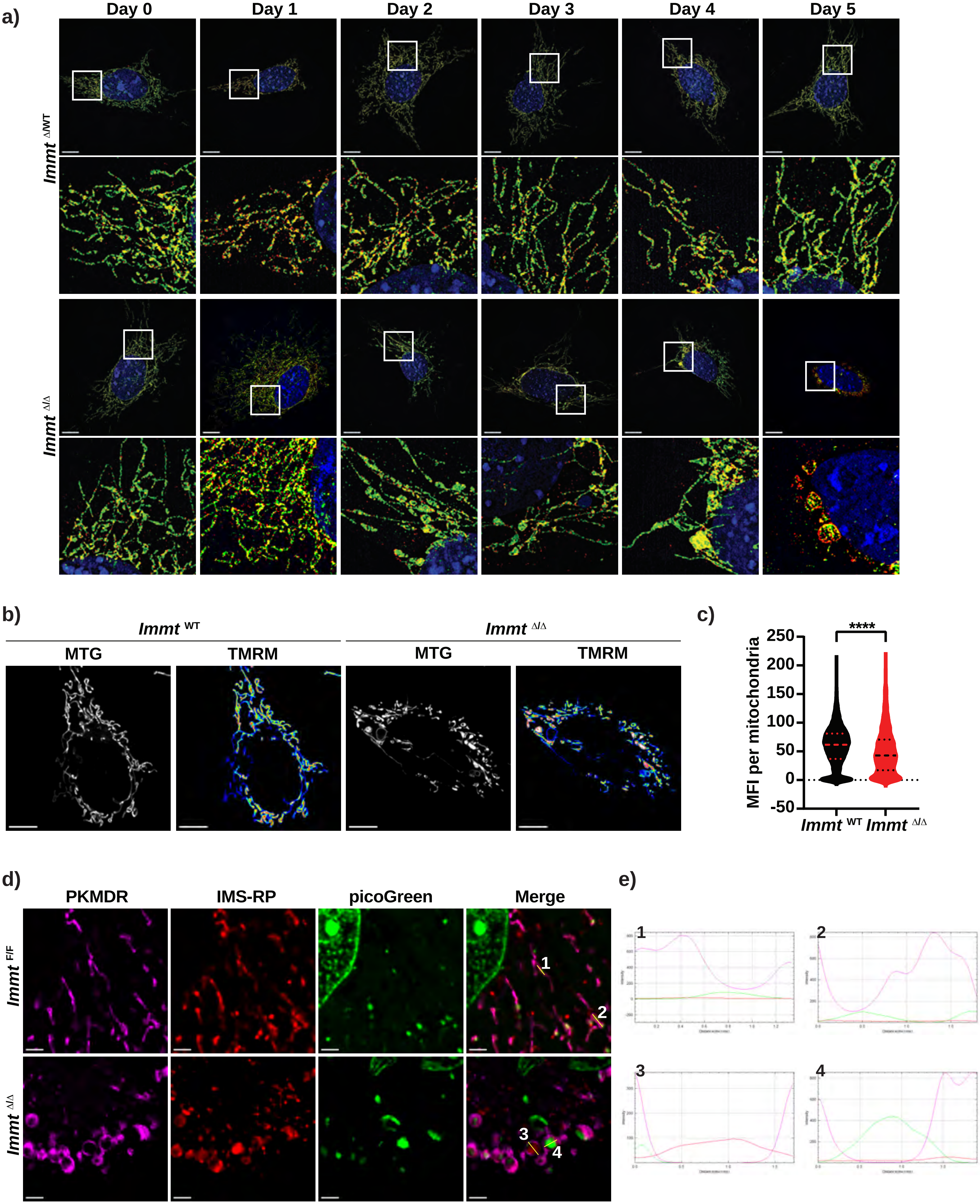

Supplemental Figure 4

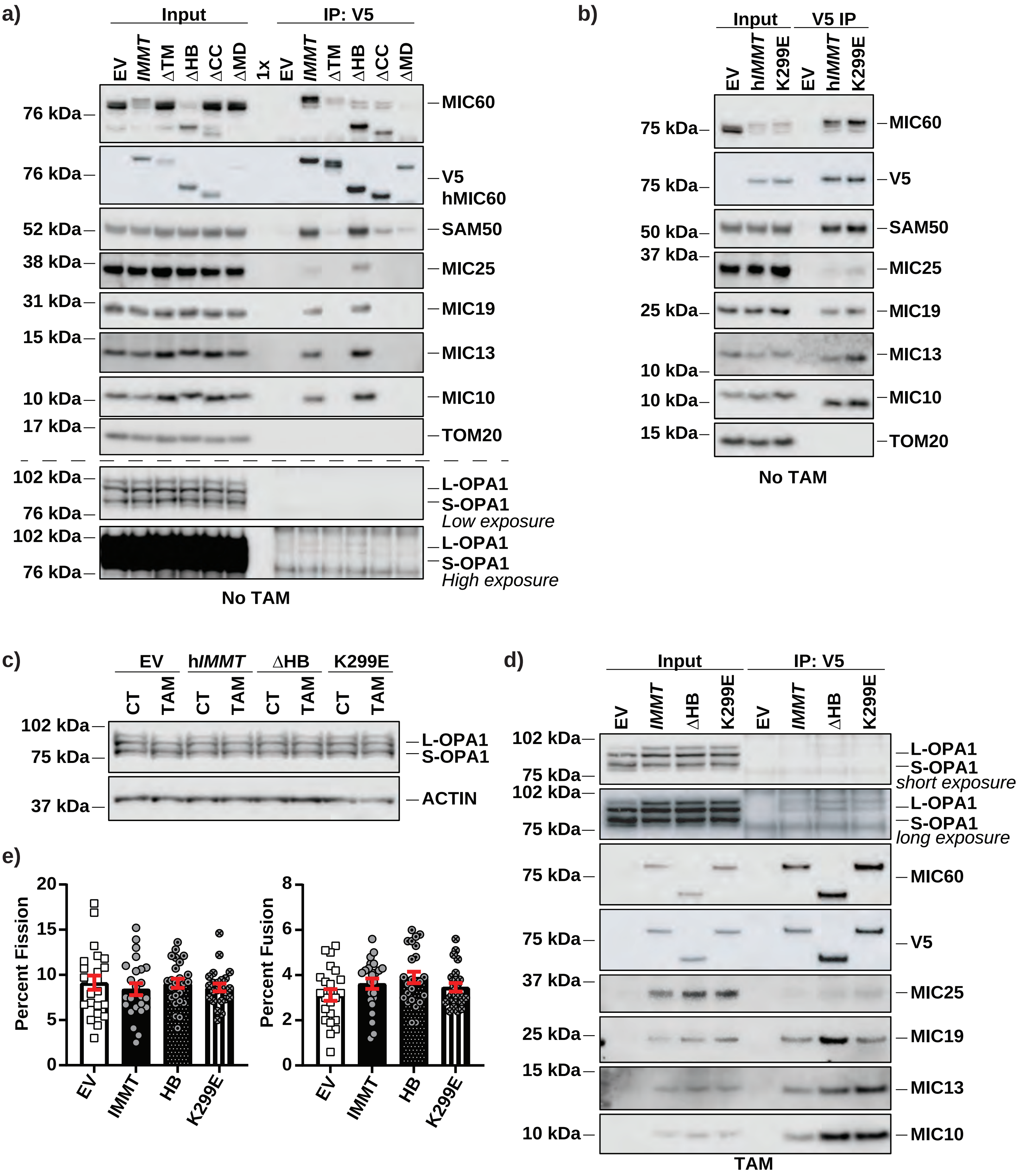

**Supplemental Tables**

**Supplemental Table 1**. Related to Figure 2. Linear mixed model (LMM) statistical analysis of morphological features, comparing *Immt*$^{f/f}$ MEFs to *Immt*$^{WT}$ MEFs after 4 days of 4-OH-tamoxifen (TAM) treatment).

| feature | Comparison | estimate | se | back_ transformed | ci_lower | ci_upper | t_value | p_value | effect_size_d | p_adj |
|---|---|---|---|---|---|---|---|---|---|---|
| solidity | Deleted:WT | 0.046 | 0.012 | 0.046 | 0.022 | 0.070 | 3.735 | 0.026 | 0.265 | 0.029 |
| circularity | Deleted:WT | 0.091 | 0.024 | 9.479 | 4.529 | 14.663 | 3.836 | 0.025 | 0.334 | 0.029 |
| area_perimeter_ratio | Deleted:WT | -0.138 | 0.042 | -12.899 | -19.750 | -5.463 | -3.304 | 0.029 | -0.294 | 0.029 |
| major_axis_length | Deleted:WT | -0.413 | 0.108 | -33.822 | -46.494 | -18.149 | -3.807 | 0.020 | -0.368 | 0.029 |
| minor_axis_length | Deleted:WT | -0.281 | 0.065 | -24.468 | -33.542 | -14.155 | -4.297 | 0.012 | -0.302 | 0.029 |

**Supplemental Table 2.** Related to Figure 2. Manual scoring of mitochondrial morphology from the live-cell tauSTED images. n=48 cells each.

| | Percent of Cells Containing: | | |
|---|---|---|---|
| | **Elongated, Branched** | **Fragmented** | **Enlarged, Rounded** |
| WT | 83.33% | 45.83% | 4.17% |
| Deleted | 60.42% | 47.92% | 91.67% |

**Supplemental Table 3**. Related to Figure 4. Linear mixed model (LMM) statistical analysis of morphological features, comparing the h*IMMT* mutant-expressing MEFs to Empty Vector (EV)-expressing *Immt*$^{f/f}$ MEFs after 4 days of 4-OH-tamoxifen (TAM) treatment.

| feature | Comparison | estimate | se | back_ transformed | ci_lower | ci_upper | t_value | p_value | effect_size_d | p_adj |
|---|---|---|---|---|---|---|---|---|---|---|
| solidity | CC:EV | 0.015 | 0.017 | 0.015 | -0.019 | 0.049 | 0.856 | 0.410 | 0.095 | 0.683 |
| solidity | HB:EV | -0.026 | 0.017 | -0.026 | -0.060 | 0.008 | -1.495 | 0.162 | -0.167 | 0.612 |
| solidity | IMMT:EV | -0.027 | 0.017 | -0.027 | -0.060 | 0.007 | -1.534 | 0.153 | -0.170 | 0.612 |
| solidity | K299E:EV | -0.026 | 0.017 | -0.026 | -0.060 | 0.008 | -1.475 | 0.168 | -0.164 | 0.612 |
| solidity | MD:EV | -0.013 | 0.017 | -0.013 | -0.047 | 0.020 | -0.784 | 0.450 | -0.086 | 0.686 |
| solidity | TM:EV | -0.028 | 0.017 | -0.028 | -0.061 | 0.006 | -1.623 | 0.133 | -0.179 | 0.612 |
| circularity | CC:EV | 0.021 | 0.018 | 2.107 | -1.400 | 5.738 | 1.169 | 0.264 | 0.083 | 0.623 |
| circularity | HB:EV | -0.026 | 0.018 | -2.547 | -5.919 | 0.946 | -1.436 | 0.174 | -0.102 | 0.612 |
| circularity | IMMT:EV | -0.041 | 0.018 | -4.061 | -7.331 | -0.674 | -2.342 | 0.037 | -0.164 | 0.612 |
| circularity | K299E:EV | -0.025 | 0.018 | -2.465 | -5.812 | 1.000 | -1.401 | 0.185 | -0.099 | 0.612 |
| circularity | MD:EV | 0.019 | 0.017 | 1.917 | -1.511 | 5.465 | 1.088 | 0.297 | 0.075 | 0.623 |
| circularity | TM:EV | 0.003 | 0.018 | 0.310 | -3.075 | 3.813 | 0.177 | 0.863 | 0.012 | 0.925 |
| area_perimeter_ratio | CC:EV | 0.067 | 0.053 | 6.966 | -3.588 | 18.675 | 1.271 | 0.224 | 0.141 | 0.612 |
| area_perimeter_ratio | HB:EV | 0.053 | 0.054 | 5.488 | -5.020 | 17.160 | 0.998 | 0.334 | 0.112 | 0.626 |
| area_perimeter_ratio | IMMT:EV | -0.021 | 0.053 | -2.059 | -11.651 | 8.575 | -0.396 | 0.699 | -0.043 | 0.873 |
| area_perimeter_ratio | K299E:EV | 0.010 | 0.053 | 0.958 | -8.999 | 12.004 | 0.180 | 0.860 | 0.020 | 0.925 |
| area_perimeter_ratio | MD:EV | -0.021 | 0.052 | -2.060 | -11.513 | 8.403 | -0.402 | 0.694 | -0.043 | 0.873 |
| area_perimeter_ratio | TM:EV | -0.054 | 0.052 | -5.304 | -14.457 | 4.829 | -1.051 | 0.311 | -0.114 | 0.623 |
| major_axis_length | CC:EV | -0.012 | 0.093 | -1.193 | -17.686 | 18.605 | -0.129 | 0.899 | -0.011 | 0.930 |
| major_axis_length | HB:EV | 0.166 | 0.094 | 18.066 | -1.784 | 41.929 | 1.768 | 0.099 | 0.157 | 0.612 |
| major_axis_length | IMMT:EV | 0.119 | 0.093 | 12.690 | -6.000 | 35.096 | 1.291 | 0.219 | 0.113 | 0.612 |
| major_axis_length | K299E:EV | 0.103 | 0.093 | 10.847 | -7.643 | 33.039 | 1.106 | 0.288 | 0.097 | 0.623 |
| major_axis_length | MD:EV | -0.032 | 0.091 | -3.197 | -19.039 | 15.744 | -0.356 | 0.727 | -0.031 | 0.873 |
| major_axis_length | TM:EV | 0.002 | 0.091 | 0.235 | -16.204 | 19.898 | 0.026 | 0.980 | 0.002 | 0.980 |
| minor_axis_length | CC:EV | 0.067 | 0.074 | 6.953 | -7.434 | 23.576 | 0.912 | 0.376 | 0.077 | 0.664 |
| minor_axis_length | HB:EV | 0.124 | 0.075 | 13.200 | -2.181 | 30.998 | 1.664 | 0.116 | 0.142 | 0.612 |
| minor_axis_length | IMMT:EV | 0.034 | 0.073 | 3.497 | -10.314 | 19.436 | 0.470 | 0.645 | 0.039 | 0.873 |
| minor_axis_length | K299E:EV | 0.056 | 0.074 | 5.779 | -8.440 | 22.207 | 0.763 | 0.458 | 0.064 | 0.686 |
| minor_axis_length | MD:EV | -0.013 | 0.072 | -1.254 | -14.255 | 13.717 | -0.175 | 0.863 | -0.014 | 0.925 |
| minor_axis_length | TM:EV | -0.043 | 0.072 | -4.170 | -16.817 | 10.400 | -0.590 | 0.564 | -0.049 | 0.806 |

St. Jude - Confidential

**Supplemental Table 4.** Related to Figure 4. Manual scoring of mitochondrial morphology from the live-cell tauSTED images. Cell numbers are as follows: EV=58, h*IMMT*=58, ΔTM=50, ΔHB=39, ΔCC=50, ΔMD=50, and K299E=50. Note that EV and h*IMMT* results combine all images from Figures 4 (n=29) and 5 (n=29).

| | Elongated, Branched | Fragmented | Enlarged, Rounded |
|---|---|---|---|
| EV | 44.83% | 65.52% | 79.31% |
| IMMT | 79.31% | 55.17% | 3.45% |
| TM | 70.00% | 52.00% | 86.00% |
| HB | 69.23% | 66.67% | 12.82% |
| CC | 16.00% | 92.00% | 88.00% |
| MD | 64.00% | 68.00% | 82.00% |
| K299E | 60.00% | 76.00% | 20.00% |

**Supplemental Table 5.** Summary of PCR primers used in this study.

| name | sequence (5'-3') |
|---|---|
| Immt-Not1-Forward | AAAAGCGGCCGCGCCACCATGCTGCGGGCCT |
| Immt-BamHI-V5-Reverse | GGGGGGATCCTCATCGAGACCGAGGAGAGGG |
| ΔTM First Fragment Reverse | CCCATTTGGCATATAGCCCAGAGCTGCCTG |
| ΔTM Second Fragment Forward | CAGGCAGCTCTGGGCTATATGCCAAATGGG |
| ΔHB First Fragment Reverse | GTTTCTCCAAGGCAACTTCTTCAGGTGG |
| ΔHB Second Fragment Forward | CCACCTGAAGAAGTTGCCTTGGAGAAAC |
| ΔCC First Fragment Reverse | CTCCAGATCACCATGCTCAATTAACGTGATGTGCTGCTTTTCGGTG |
| ΔCC Second Fragment Forward | CACCGAAAAGCAGCACATCACGTTAATTGAGCATGGTGATCTGGAG |
| ΔMD-V5 Reverse | GGGGGATCCTCATCGAGACCGAGGAGAGGGTTAGGGATAGGCTTACCAACC<br>ACTTTGTACAAGAAAGTTGGGATGCAATAGGAAGC |
| K299E SDM Forward | GATGCCCTTCTCAAAGCCGAAGAAGAGTTAGAGAAGA |
| K299E SDM Reverse | TCTTCTCTAACTCTTCTTCGGCTTTGAGAAGGGCATC |

St. Jude - Confidential

**Supplemental Figure Legends**

**Supplemental Figure 1. *Immt* deletion in liver does not induce apoptosis. a)** Western blot analysis of whole liver lysates derived from adult wild-type (WT) and *Immt*$^{\Delta/\Delta}$ mice across 5 after LP1-Cre injection. Mouse embryonic fibroblasts (MEFs) treated with staurosporine (STS) for 24 h were used as a control for caspase-3 cleavage. Data is representative of 3 independent experiments. **b)** Western blot analysis of liver, spleen, muscle, and heart tissues lysates derived from adult WT and *Immt*$^{\Delta/\Delta}$ mice 3 weeks after LP1-Cre injection (n = 3 mice/group, representative of 3 independent experiments). **c)** Western blot analysis of whole liver lysates derived from adult WT and *Immt*$^{\Delta/\Delta}$ mice 3 weeks after LP1-Cre injection, assessing markers of the electron transport chain (n = 3 mice/group, representative of 3 independent experiments). # indicates mitochondrial-encoded proteins. **d)** Representative histology images of caspase-3 and TUNEL in whole liver lysates derived from adult wild-type (WT; n = 4 M, 3 F) and *Immt*$^{\Delta/\Delta}$ (n = 4 M, 5 F) mice 3 weeks after LP1-Cre injection. Scale bar = 100 µm **e)** Changes in seral liver damage markers alanine transaminase (ALT) and aspartate aminotransferase (AST) after LP1-Cre injection in WT (n = 4 M, 6 F) and *Immt*$^{\Delta/\Delta}$ (n = 7 M, 2 F) mice. Data is presented as mean ± SEM from 3 independent experiments. **f)** Western blot analysis of cytochrome *c* release from isolated liver mitochondria 3 weeks after LP1-Cre injection. Mitochondria were treated with the BID–stapled alpha-helical peptide (SAHB). S = supernatant, P = pellet. For all panels, data is representative of at least 3 independent experiments.

**Supplemental Figure 2. *Immt* deletion *in vitro* does not induce apoptosis. a)** Western blot analysis of control (heterozygous *Immt*$^{\Delta/WT}$) and *Immt*$^{\Delta/\Delta}$ mouse embryonic fibroblasts (MEFs) over 5 days after treatment with 4-OH-tamoxifen (TAM) or DMSO (control, Day 0). Data is representative of 3 independent experiments. **b)** Agilent Seahorse analysis for mitochondrial oxygen consumption (ATP production, left; coupling efficiency, middle; proton leak, right) in heterozygous *Immt*$^{\Delta/WT}$ and *Immt*$^{\Delta/\Delta}$ MEFs over 5 days treatment with TAM. Results depict the fold change of TAM to control (DMSO, Day 0) and represent 3 independent experiments. **c)** Agilent Seahorse analysis for mitochondrial oxygen consumption in WT and *Immt*$^{\Delta/\Delta}$ MEFs 4 days post-treatment with TAM. Results depict the fold change of TAM to control (DMSO) and represent 3 independent experiments. **d)** MEF proliferation across 7 days after control or TAM treatment. **e)** Viability of MEFs on day 7 of treatment. Results are shown as a fold change of TAM to control. **f)** Western blot analysis of MEFs on day 4 of control or TAM treatment, with MEFs treated with staurosporine (STS) for 24 h serving as a control for caspase-3 cleavage. For all panels, data represents 3 independent experiments; data is presented as the mean ± SEM in (b–d). Statistical significance was determined by one-way ANOVA (b) or an unpaired Welch's *t*-test (c–e). * $p \leq 0.05$, ** $p \leq 0.01$, *** $p \leq 0.001$, **** $p \leq 0.0001$, ns = no significance.

**Supplemental Figure 3. *In vitro Immt* deletion reveals intermembrane space–filled regions and differential membrane potential. a)** Representative structured illumination microscopy images of wild-type (WT; *Immt*$^{\Delta/WT}$) and *Immt*$^{\Delta/\Delta}$ mouse embryonic fibroblasts (MEFs) across 5 days after 4-OH-tamoxifen (TAM) treatment. MEFs were stained with DAPI (blue), TOM20 (red), and cytochrome *c* (green). Images represent 3 independent experiments, scale bar = 10 µm. **b)** Mitochondrial membrane potential in WT or *Immt*$^{\Delta/\Delta}$ MEFs 4 days post TAM treatment. Cells were stained with MitoTracker Green (MTG, gray) and TMRM (multi-colored based on signal intensity). Images represent 3 independent experiments, scale bar = 10 µm. **c)** Violin plot depicting mean fluorescence intensity (MFI) of the TMRM signal per mitochondria in WT (n=88) or *Immt*$^{\Delta/\Delta}$

St. Jude - Confidential

(n=81) MEFs after 4 days of TAM treatment. Statistical significance determined using an unpaired Welch's *t*-test; **** $p \leq 0.0001$. **d)** Representative images of MEFs expressing IMS-RP (red) and treated with DMSO control (*Immt*$^{f/f}$, n=10) or TAM (*Immt*$^{\Delta/\Delta}$, n=10) for 4 days. Cells were stained with picoGreen (green) and PKMito-Deep Red (magenta). Scale bar = 2 µm. **e)** Line analysis of selected mitochondria from (d).

**Supplemental Figure 4. Mitochondrial dynamics are unchanged in mouse embryonic fibroblasts with h*IMMT* domain mutants. a)** Western blot analysis of immunoprecipitated proteins pulled down with V5 from mouse embryonic fibroblasts (MEFs) expressing h*IMMT* domain mutants (transmembrane domain [ΔTM], helical bundle [ΔHB], the coiled-coil helix [ΔCC], and mitofilin domain [ΔMD]) after 4 days DMSO (control; CT) treatment. **b)** Western blot analysis of immunoprecipitated proteins pulled down with V5 from MEFs expressing an empty vector (EV), h*IMMT*, or K299E mutant after 4 days CT treatment. **c)** Western blot analysis of OPA1 in MEFs stably expressing EV, h*IMMT*, ΔHB, or K299E and treated for 4 days with CT or 4-OH-tamoxifen (TAM). **d)** Western blot analysis of immunoprecipitated proteins pulled down with V5 from MEFs expressing EV, h*IMMT*, ΔHB, or K299E mutant after 4 days of TAM treatment. **e)** Differences in fission and fusion dynamics in MEFs expressing EV, h*IMMT*, ΔHB, or K299E after 4 days of TAM treatment (n=23 for each cell group). Data is presented as the mean ± SEM from 3 independent experiments. For all western blots, data is representative of 3 independent experiments.

**Supplemental Video 1.** Related to Figure 1f. Three-dimensional rotation of the focused ion beam electron microscopy stack of mitochondria isolated from LP1-Cre injected into WT mice.

St. Jude - Confidential

**Supplemental Video 2.** Related to Figure 1f. Three-dimensional rotation of the focused ion beam electron microscopy stack of mitochondria isolated from LP1-Cre injected into *Immt*$^{f/f}$ mice.

St. Jude - Confidential
