## Supplementary figures and images for "Protein domain characterization reveals human MIC60 tolerates loss of helical bundle domain"

### Supplemental Videos

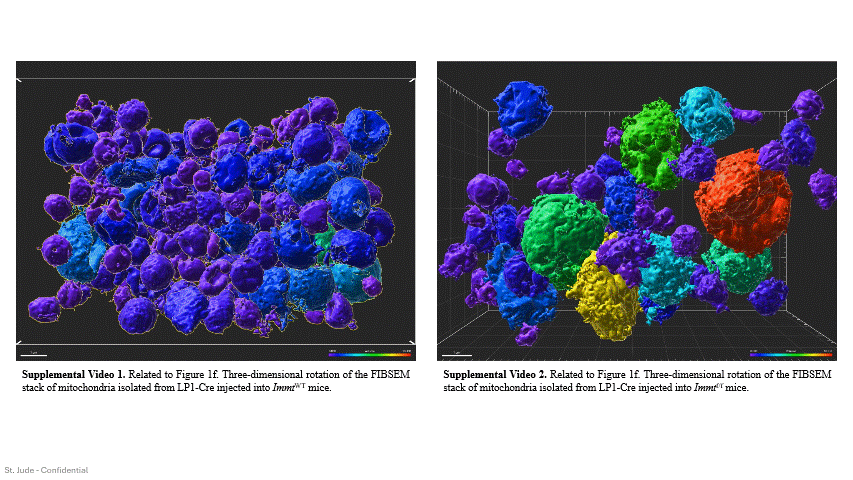
